# How enhancers regulate wavelike gene expression patterns: Novel enhancer prediction and live reporter systems identify an enhancer associated with the arrest of pair-rule waves in the short-germ beetle *Tribolium*

**DOI:** 10.1101/2022.09.09.507237

**Authors:** Christine Mau, Heike Rudolf, Frederic Strobl, Benjamin Schmid, Timo Regensburger, Ralf Palmisano, Ernst Stelzer, Leila Taher, Ezzat El-Sherif

## Abstract

A key problem in development is to understand how genes turn on or off at the right place and right time during embryogenesis. Such decisions are made by non-coding sequences called ‘enhancers’. Much of our models of how enhancers work rely on the assumption that genes are activated *de novo* as stable domains across embryonic tissues. Such view has been strengthened by the intensive landmark studies of the early patterning of the anterior-posterior (AP) axis of the *Drosophila* embryo, where indeed gene expression domains seem to arise more or less stably. However, careful analysis of gene expressions in other model systems (including the AP patterning in vertebrates and short-germ insects like the beetle *Tribolium castaneum*) painted a different, very dynamic view of gene regulation, where genes are oftentimes expressed in a wavelike fashion. How such gene expression waves are mediated at the enhancer level is so far unclear. Here we establish the AP patterning of the short-germ beetle *Tribolium* as a model system to study dynamic and temporal pattern formation at the enhancer level. To that end, we established an enhancer prediction system in *Tribolium* based on time- and tissue-specific ATAC-seq and an enhancer live reporter system based on MS2 tagging. Using this experimental framework, we discovered several *Tribolium* enhancers, and assessed the spatiotemporal activities of some of them in live embryos. We found our data consistent with a model in which the timing of gene expression during embryonic pattern formation is mediated by a balancing act between enhancers that induce rapid changes in gene expressions (that we call ‘dynamic enhancers’) and enhancers that stabilizes gene expressions (that we call ‘static enhancers’).

## Introduction

While an embryo is growing, each cell continuously receives inputs from surrounding cells. The cell processes these inputs and decides its fate accordingly. This decision-making process relies on non-coding sequences called ‘enhancers’ (1,2). Much of our models of how enhancers work during development relies on the assumption that genes are activated *de novo* across embryonic tissues as stable domains of gene expression (3–5), that then undergo little or no change, either indefinitely or until they do their job whereafter they gradually fade away. Such a view has been strengthened by the intensive landmark studies of the early patterning of the anterior-posterior (AP) axis of the fruit fly *Drosophila melanogaster* embryo, where indeed gene expression domains seem to arise more or less stably (reviewed in (6–9)). However, careful analysis of gene expressions in other model systems painted a different, very dynamic view of gene regulation (9,10). For example, during the AP patterning of vertebrates, oscillatory waves of gene expressions were shown to sweep the embryo before they stabilize into their final positions (9,11–16), demarcating future vertebrae. Likewise, Hox gene expressions propagate along the AP axis of vertebrates, demarcating future axial identities (9,17–21). In both neural tube and limb bud of vertebrates, gene expressions arise in a temporal sequence and spread across the tissue, dividing them into different embryonic fates (22–27).

Surprisingly, patterning of the AP axis of insects, the same process that popularized the static view of gene regulation, turned out to be much more dynamic than previously thought. In insects, the AP axis is divided into segments via the striped expression of a group of genes called ‘pair-rule’ genes, and into domains of different axial fates via the expression of a group of genes called ‘gap genes’ (6,9). In the flour beetle *Tribolium castaneum*, a short-germ insect thought to adopt a more ancestral mode of AP patterning than long-germ insects like *Drosophila*, both pair-rule and gap genes are expressed as dynamic waves that propagate from posterior to anterior (28–32). Similar dynamics seem to be involved in segmentation in other insects and arthropods (33–39). Even pair-rule and gap genes in *Drosophila*, classically thought to be expressed stably, were shown more recently to undergo dynamic (albeit limited) posterior-to-anterior shifts (40–44), a phenomenon that has been suggested to be an evolutionary vestige of outright gene expression waves of the sort observed in *Tribolium* (9,45–48). These observations show that the static view of gene regulation, once popularized by classical studies of AP patterning in *Drosophila*, is inaccurate and that gene regulation is in most cases a dynamic phenomenon. Hence, new models of embryonic pattern formation – and concomitantly, new models of how enhancers work within pattern formation models – are needed.

Some of the authors have recently suggested a model that explains the generation of either periodic or non-periodic waves of gene expressions, termed the ‘Speed Regulation’ model (**Figure 1A,B**) (9,31,32,45,49). In this model, a morphogen gradient (of a molecular factor we termed the ‘speed regulator’) modulates the speed of either a molecular clock or a genetic cascade. This scheme was shown (*in silico*) to be able to produce periodic waves in the former case (**Figure 1A**), and non-periodic waves in the latter (**Figure 1B**). The model was shown to be involved in generating pair-rule and gap gene expression waves in the early *Tribolium* embryo (30,31), and consistent with recent findings in vertebrate somitogenesis (50–52). Furthermore, a molecular model, termed the ‘Enhancer Switching’ model (**Figure 1C,D**), has been suggested as a mechanism for how a morphogen gradient could fine-tune the speed of a clock or a genetic cascade, serving as a molecular realization of the Speed Regulation model (9,31,49). The Enhancer Switching model posits that each patterning gene is simultaneously wired into two gene regulatory networks (GRNs) (**Figure 1C**): (*i*) a dynamic GRN that drives periodic or sequential gene activities, and (*ii*) a static GRN that stabilizes gene expressions. The concentration of the speed regulator (shown in gray in **Figure 1C**) activates the dynamic GRN while represses the static GRN, and hence sets the balance between the contribution of each GRN to the overall dynamics and, consequently, the speed of gene regulation (9,31,49). At high concentrations of the speed regulator, the dynamic GRN is more dominant than the static one, and hence fast oscillations or sequential gene activities are mediated. On the other hand, at low concentrations of the speed regulator, the static GRN is more dominant, and hence slow oscillations or sequential gene activities are mediated. As mentioned, the model posits that each gene is wired into two different GRNs, a requirement that was suggested to be realized using two enhancers per patterning gene: (*i*) a dynamic enhancer that encodes the wiring of the gene within the dynamic GRN, and (*ii*) a static enhancer that encodes the wiring of the gene within the static GRN (**Figure 1D**). The model is partially supported (or rather inspired) by observations in the early *Drosophila* embryo, where the gap gene *Krüppel* (*Kr*) was shown to be regulated by two enhancers whose activities resemble those predicted by the Enhancer Switching model (31,41). Similar observations were made for the *Drosophila* gap gene *giant* (*gt*) (53). Furthermore, in vertebrates, it has been suggested that some enhancers or genetic programs mediate the initiation of segmentation clock waves posteriorly, and others mediate their anterior expressions (54–56).

**Figure 1.**
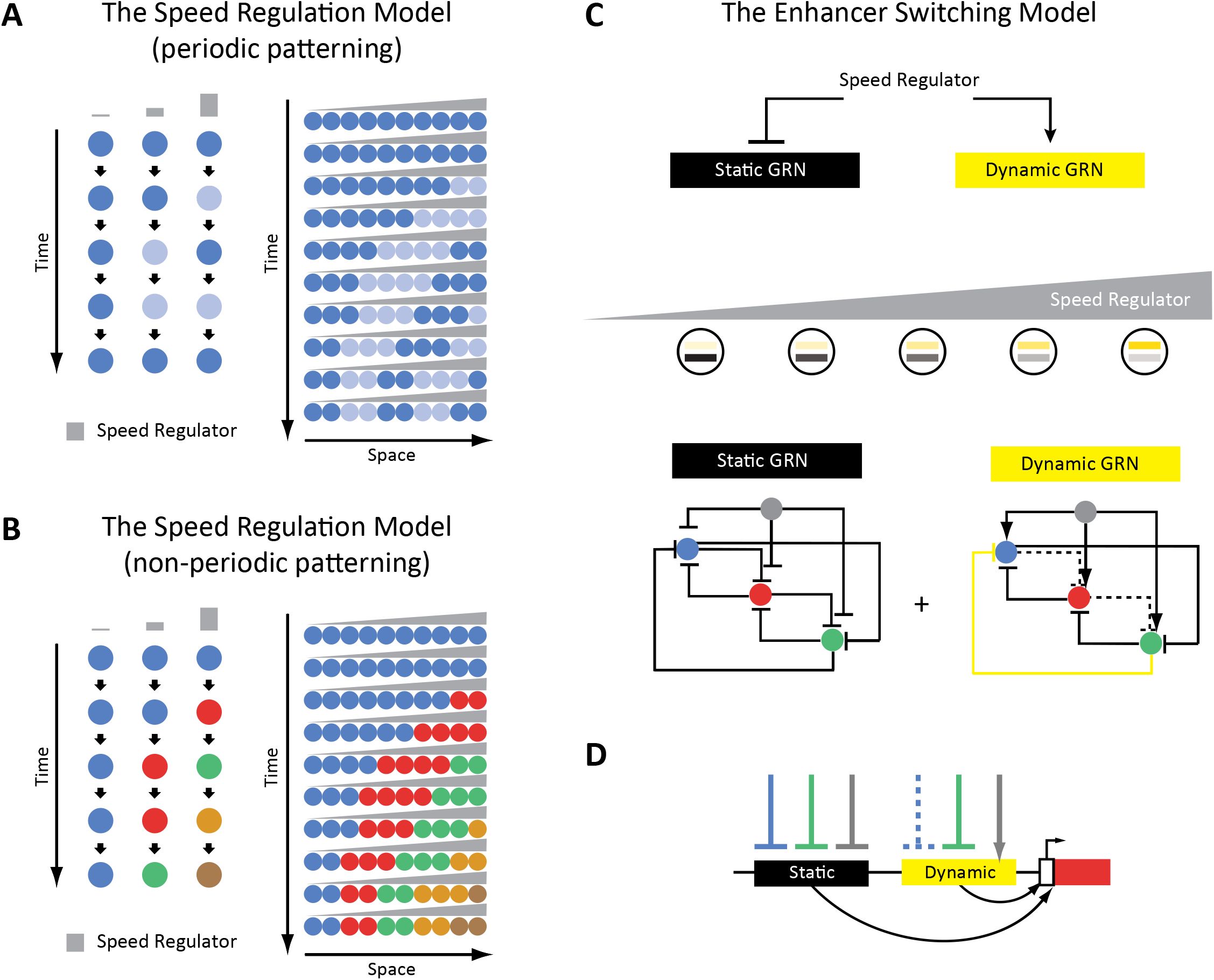
The Enhancer Switching Model as a molecular realization of the Speed Regulation Model. (**A**) The Speed Regulation Model for periodic patterning. Left: Cells can oscillate between two states mediated by a molecular clock: high (shown in dark blue), and low (shown in light blue). The concentration of the speed regulator (shown in gray) modulates the speed of the molecular clock (i.e. its frequency). Right: A gradient of the speed regulator across a tissue (represented by a row of cells) induces a periodic wave that propagates from the high to the low end of the gradient. (**B**) The Speed Regulation Model for non-periodic patterning. Same as for the periodic case (A), except the molecular clock is replaced with a genetic cascade that mediates the sequential activation of cellular states (represented by different colors). (**C**) The Enhancer Switching Model, a molecular realization of the Speed Regulation model, is composed of two GRNs: one dynamic and one static. The dynamic GRN can be either a clock (to mediate periodic patterning) or a genetic cascade (to mediate non-periodic patterning). The static GRN is a multi-stable gene circuit that mediates the stabilization of gene expression patterns. The speed regulator activates the dynamic GRN but represses the static GRN, and so a gradient of the speed regulator (shown in gray) mediates a gradual switch from the dynamic to the static GRN along the gradient. Shown example realizations of dynamic and static GRNs, where the dynamic GRN represents either a molecular clock or a genetic cascade, depending on the absence or presence of the repressive interaction shown in yellow, respectively. (**D**) Separate dynamic and static enhancers encode the wiring of each gene (shown here only for the red gene) within the dynamic and static GRNs, respectively.

Most of enhancers regulating AP patterning have been discovered and characterized in *Drosophila*, and so early patterning of the *Drosophila* embryo might seem like a good model system to study enhancer regulation of dynamic gene expressions (57,58). However, the observed gene expression dynamics of *Drosophila* gap and pair-rule genes are only vestigial, not outright waves of the sort observed during the AP patterning of vertebrates or short-germ insects like *Tribolium*. Hence, we sought to test the predictions of our Enhancer Switching model in a system where ‘canonical’ gene expression waves are observed. We thought that the AP patterning of *Tribolium* serves our purpose well, and more generally, is an excellent model system to study enhancer regulation of dynamic gene expressions. First, *Tribolium* exhibits robust systemic RNAi, which greatly eases the generation of RNAi knockdowns using parental RNAi (59–62). Second, AP patterning takes place in the early *Tribolium* embryo, which eases the interpretation of RNAi knockdowns generated using parental RNAi, without the need of time-specific or tissue-specific genetic perturbations. Indeed, most genetic interactions between gap and pair-rule genes in *Tribolium* have been elucidated by a handful of labs over a few years since the discovery of the efficacy of parental RNAi in *Tribolium* (31,32,63–74). Third, several genetic and genomic tools have been developed for *Tribolium* (75): transgenesis (76,77), transposon-based random mutagenesis (78), large-scale RNAi screens (62), CRISPR-Cas9 (79,80), live imaging (29,81–83), and a tissue culture assay (29,84). Fourth, the *Tribolium* genome is compact compared to vertebrate model systems, rendering enhancer discovery more tractable in that organism.

Thus, in this work, we sought to establish the patterning of the early *Tribolium* embryo as a model system for studying enhancer regulation of dynamic gene expressions and wavelike gene expression patterns. To that end, we set to (*i*) discover enhancer regions that regulate early patterning genes in *Tribolium*, and (*ii*) characterize the spatiotemporal activity dynamics of these enhancers.

Several strategies can be used to predict enhancer regions, each with their own advantages and disadvantages. Assaying open chromatin is a popular method. In particular, “Assay for Transposase-Accessible Chromatin with high-throughput sequencing” (ATAC-seq) (85) is fast and sensitive, and requires very little embryonic tissue (often one embryo, or even a tissue dissected from one embryo) compared to other open chromatin assays. Nevertheless, not all open chromatin regions are active enhancers. Chromatin is also accessible at promoters, insulators, and regions bound by repressors (86–89), and hence, enhancer discovery using open chromatin assays has a high false positive rate. Interestingly, chromatin accessibility has been shown to be dynamic across space and time at active developmental enhancers compared to other regulatory elements like promoters (90–92), and therefore, dynamic chromatin accessibility has been proposed as an accurate predictor for active enhancers. Thus, in this paper, we used a time-specific and tissue-specific ATAC-seq assay to elucidate the dynamics of open chromatin in space and time in the early *Tribolium* embryos, used the assay to discover a number of active *Tribolium* enhancers, and assessed the association between differential ATAC-seq peak accessibility and enhancer activity.

The second step to understand how enhancers mediate dynamic gene expressions and wavelike gene expression patterns is to characterize the spatiotemporal dynamics driven by the discovered enhancers. *In situ* staining of carefully staged embryos can go a long way in characterizing dynamic gene expressions. However, salient features of these dynamics can be missed using this method, and a strategy to visualize enhancer activities in live embryos is thus needed. Using fluorescent proteins (FP) as reporters for enhancer activities has been traditionally the method of choice in live imaging studies. Nonetheless, FPs suffer from low degradation rates, which results in averaging out of fast changing gene expressions, rendering them unsuitable for visualizing highly dynamic gene activities. This problem can be circumvented by using a destabilized version of FPs (93). However, the reported reduction in FP, although decent, is not enough to detect fast dynamics of many developmental genes. Moreover, destabilized FP suffers from low fluorescent intensity (94). Another strategy is to tag RNAs (95), like in MS2 tagging (96), where MS2 tandem repeats are inserted within a reporter gene. Upon reporter gene activation, the MS2 repeats are transcribed into stem loops that readily bind MS2 virus coat protein (MCP). If MCP-FP fusion proteins are ubiquitously present in the background, they are then recruited at the transcription site in as many numbers as RNA polymerases are actively transcribing the MS2 reporter gene, offering a natural form of signal amplification. This strategy can be used to visualize *de novo* transcription (41,42,97–99), avoiding the averaging effect of using FPs as reporters. Therefore, to study the dynamics of gene expression waves during embryogenesis, we established an MS2-tagging system in *Tribolium*, and used it to visualize the activities of some of the enhancers we discovered using our enhancer discovery system.

In summary, we established in this paper a framework for enhancer discovery and enhancer activity visualization in both fixed and live embryos in *Tribolium*. First, we assayed the dynamics of open chromatin in space and time in the *Tribolium* embryo using ATAC-seq, and used the assay to discover a number of active enhancers. Of importance to future efforts in that vein, we found that active enhancer regions overlap with chromatin accessible sites that significantly vary across the AP axis of the embryo. Second, we established an MS2-MCP enhancer reporter system in *Tribolium* to visualize the activity dynamics of discovered enhancers in both fixed and live embryos. Using this enhancer reporter system, we showed that some of the discovered enhancers regulating gap and pair-rule genes feature expression patterns that are in line with the Enhancer Switching model.

## Results

### Profiling chromatin accessibility landscape along the AP axis of the early *Tribolium* embryo

Genomic regions of increased chromatin accessibility are typically endowed with regulatory activity (92,100). At enhancers in particular, chromatin accessibility has been shown to be dynamic across space and time, and so we set to assay the dynamics of the accessible chromatin landscape in the *Tribolium* embryo. To that end, we dissected the *Tribolium* embryo at the germband stage into three regions across its AP axis (**Figure 2A**): anterior (‘a’), middle (‘m’), and posterior (‘p’), and performed ATAC-seq on each region. We did that for two time points: 23-26 hours after egg lay (AEL) (hereafter, termed IT23), and 26-29 hours AEL (hereafter, termed IT26) (**Materials and Methods**). Our experimental design included six sample groups (3 regions x 2 time points) with 2-3 biological replicates. We generated and sequenced 17 ATAC-seq libraries to an average depth of 1,835,762 unique, high-quality pairs of reads (3.6X genomic coverage, **Materials and Methods**). Biological replicates of our ATAC-seq libraries were highly similar with a median Spearman’s correlation coefficient of 0.875 (**Supplementary Figure 1**), demonstrating the reproducibility of the data.

**Figure 2.**
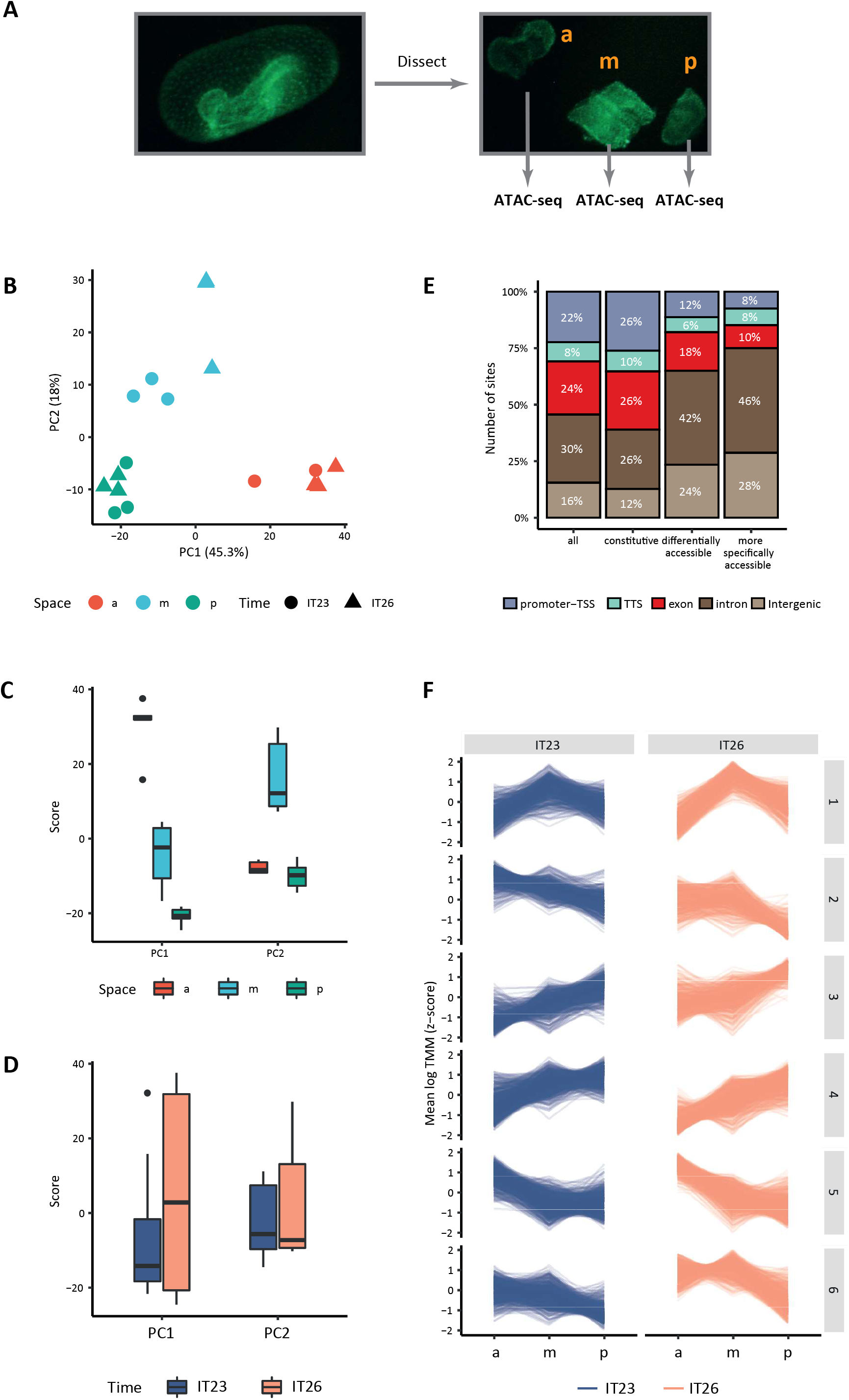
Chromatin accessibility dynamics during AP patterning of the early *Tribolium* embryo. (**A**) Embryos 23-26 h AEL (IT23) or 26-29 h AEL (IT26) were dissected and cut into anterior (a), middle (m), and posterior (p) part. Left: nGFP embryo in the eggshell. Right: dissected and cut nGFP embryo. (**B**) Principal component analysis (PCA) on the accessibility scores of the most highly accessible and variable sites in the dataset. PCA was performed using the PCA() function in the “FactoMineR” R package with default parameters. Only the first two principal components (PC) of the data are represented. The first PC explains 45.3% of the variance in the data, the second PC, 18.0%. (**C**, **D**) Boxplots showing the scores of PC1 and PC2 by space (C) and time (D). The thick line indicates the median (2nd quartile), while the box represents interquartile range (IQR, 1st to 3rd quartiles). Outliers are shown as dots. (**E**) Genomic annotation of different classes of chromatin accessible sites: all (consensus) sites, constitutive sites (i.e., consensus sites that are not differentially accessible), all differentially accessible sites, and most specifically accessible sites (i.e., sites differentially accessible in four or more comparisons). (**F**) K-means clustering of accessibility scores for differentially accessible sites. Accessibility scores have been z-score scaled for each site.

We identified a total of 12,069 chromatin accessible sites (**Materials and Methods**), with 4,017 being specific to one or two particular regions of the germband and 1,610 to a given time point (**Supplementary Figure 2**). In agreement with the ability of ATAC-seq to detect distal regulatory elements in the genome (85), a large proportion of these sites were intergenic or intronic (46%, **Supplementary Figure 3**). Principal component analysis (PCA) of most variable accessible sites (**Materials and Methods**) mainly separated the samples along the AP axis of the embryo (**Figure 2B–D**), where the first principal component (PC1) accounted for 45.3% of the variance, whereas PC2 (18.0% of the variance) predominantly distinguished the middle of the germband from its anterior and posterior ends. Among all 12,069 sites, 3,106 (26% of the accessible genome) were differentially accessible when compared between different regions along the AP axis and/or time point (**Materials and Methods**). For 1,049 of those sites, changes in accessibility were observed in four or more comparisons, indicating more intricate and, generally, specific patterns of accessibility (**Supplementary Figure 4**). Remarkably, while 62% of the constitutively accessible sites corresponded to promoters and gene bodies, 66% of the differentially accessible sites were intergenic or intronic, and this proportion was even higher (74%) among the differentially accessible sites with more specific patterns of accessibility (**Figure 2E**), suggesting that spatiotemporal control of transcription in the early *Tribolium* embryo is largely mediated by enhancers as opposed to promoters. Accessible sites were enriched for binding sites of several transcription factors (**Materials and Methods**; **Supplementary Figure 5**). In particular, motifs consistent with the binding sites of 20 different transcription factors (Abd-A, Abd-B, Ap, Awh, Cad, CG18599, CG4328, E5, Ems, Eve, Ind, Lab, Lbe, Lbl, Lim3, Pb, PHDP, Pho, Vfl, and Zen) were enriched among all types of sites, independently of their accessibility dynamics.

When comparing accessibility along the AP axis at a particular time point, 2,109 sites were differentially accessible, the majority of them along the AP axis at IT26. In addition, only 132 sites were differentially accessible between IT26 and IT23 at the same portion of the embryo (**Supplementary Figure 4B and 4C**). To gain a better insight into the spatial and temporal dynamics of chromatin accessibility, we clustered all differentially accessible sites across the germband regions and time points (**Figure 2F**). Almost half of the sites were either not accessible in the anterior region of the embryo but accessible in the middle and posterior regions (cluster 4,756 sites), or not accessible in the middle and posterior regions of the embryo but accessible in the anterior region (cluster 5,698 sites). Approximately 25% of the sites showed predominantly monotonic changes in accessibility along the AP axis at IT23, either increasing (clusters 3,420 sites) or decreasing (cluster 2,347 sites) from the anterior towards the posterior region of the embryo. Fifteen-percent of the sites showed similar accessibility levels in the anterior and middle part of the embryo, and decreased accessibility in the anterior part of the embryo (cluster 6,477 sites). Finally, 13% of the sites were most accessible in the middle region of the embryo (cluster 1,408 sites). The sites exhibited similar trends along the AP axis at both time points, although with the exception of the sites in clusters 2 and 4, the sites were generally more accessible at IT26 than at IT23. Functional enrichment analysis of the gene nearest to each site (**Materials and Methods**) revealed that while all clusters are associated with “developmental process” and “anatomical structure development”, clusters 1 and 3 are specifically related to “pattern specification process” and “regionalization” and cluster 3 is specifically associated with “anterior/posterior pattern specification” (**Supplementary Figure 6**).

Together, our findings indicate that changes in chromatin accessibility in *Tribolium* at this developmental stage are primarily associated with space rather than time, and are particularly evident when comparing the anterior part of the germband to the middle and posterior parts. Furthermore, our data suggests that sites for which accessibility varies across space are especially likely to be associated with enhancer activity, laying the foundation for a promising enhancer prediction strategy based on differential ATAC-seq peak analysis. Before assessing this proposition, however, we set to establish an enhancer reporter system to validate the activity of predicted enhancers, and analyze their transcriptional dynamics.

### Establishing an MS2-MCP enhancer reporter system to visualize enhancer activity in fixed and live *Tribolium* embryos

An enhancer reporter system has been previously established in *Tribolium* using *mCherry* as a reporter gene (101). However, long half lives of *mCherry* mRNA and proteins could average out fast transcriptional dynamics, precluding the analysis of gene expression waves. To circumvent this, we created a *Tribolium* enhancer reporter system capable of visualizing *de novo* transcription in both fixed and live embryos. Our enhancer reporter system is composed of the gene *yellow*, which has a long intron (2.7 kb). Visualizing intronic transcription of the reporter gene *yellow* using *in situ* staining in fixed embryos enables the detection of *de novo* transcription, and has been routinely used to analyze fast transcriptional dynamics in enhancer reporter experiments in *Drosophila* (102). To visualize *de novo* transcription in live *Tribolium* embryos, we set to (*i*) modify the *yellow* reporter gene to allow for MS2 tagging, and (*ii*) create a *Tribolium* transgenic line with ubiquitous expression of an MCP-FP fusion. To that end, we created two piggyBac reporter constructs: ‘enhancer>MS2-*yellow*’ (**Figure 3A**) and ‘aTub>MCP-mEmerald’ (**Figure 3B**). For the enhancer>MS2-*yellow* construct, we place an enhancer of interest upstream of the *Drosophila* Synthetic Core Promoter (DSCP) and a MS2-*yellow* reporter gene. The MS2-*yellow* reporter is composed of 24 tandem repeats of MS2 stem loops inserted in the 5’ UTR of the *yellow* gene, followed by an SV40 poly(A) tail. For the aTub>MCP-mEmerald line, we created a piggyBac construct in which the ubiquitous alpha-tubulin promoter was placed upstream of a nuclear localization sequence (NLS) and an MCP-mEmerald fusion, followed by an SV40 poly(A) tail.

**Figure 3.**
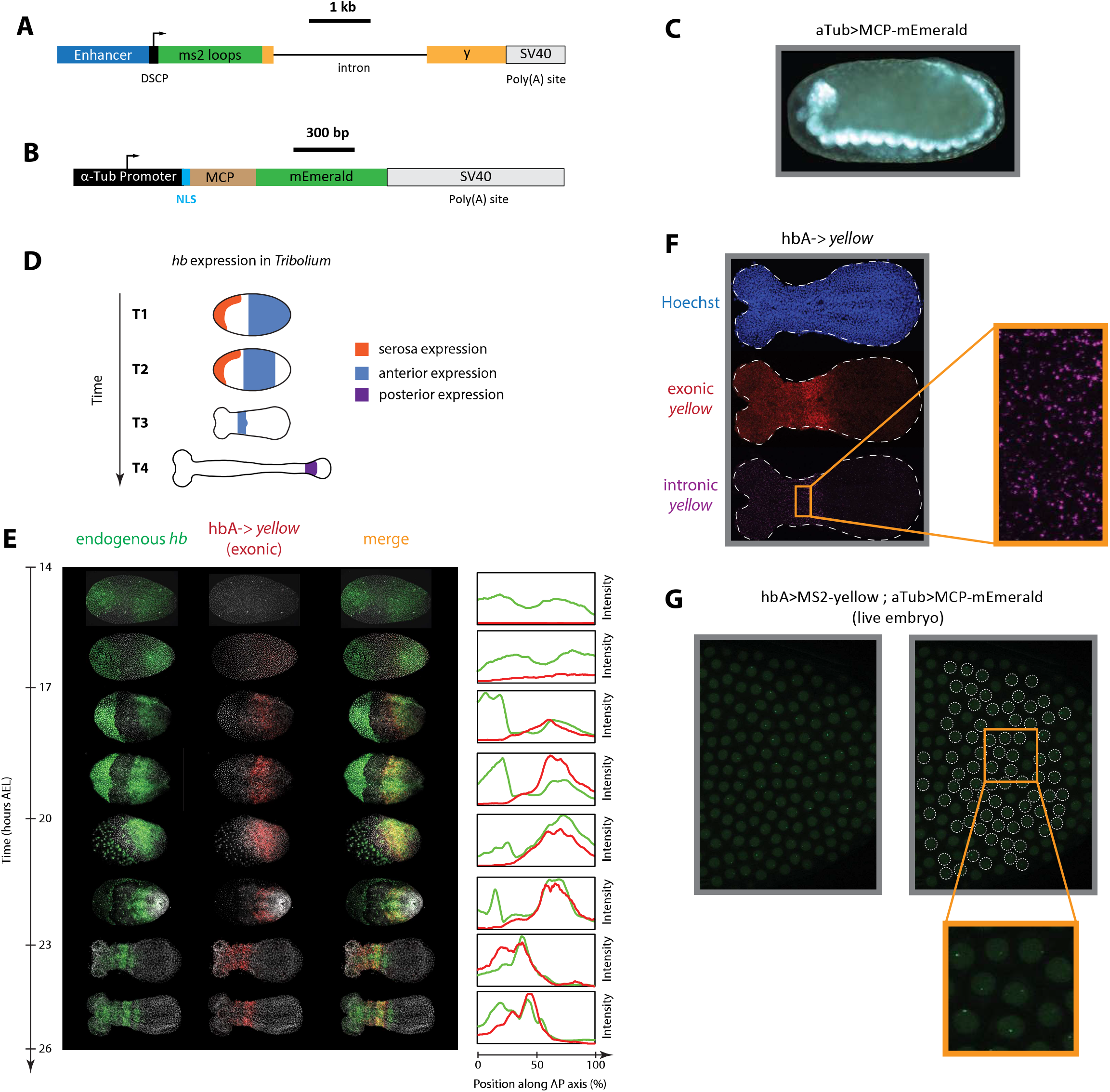
An MS2-MCP enhancer reporter system to visualize enhancer activity in fixed and live *Tribolium* embryos tested using the *Tribolium hb* enhancer hbA. (**A**) Our enhancer reporter construct: An enhancer of interest is placed upstream of a DSCP, followed by 24 tandem repeats of MS2 stem loops, the gene *yellow*, then an SV40 poly(A) tail. **(B)** The aTub>MCP-mEmerald construct: ubiquitous alpha-tubulin promoter was placed upstream of an NLS and an MCP-mEmerald fusion, followed by an SV40 poly(A) tail. (**C**) Overview image of an aTub>MCP-mEmerald embryo at the germband stage. (**D**) A schematic showing *hb* expression in *Tribolium*. In the early blastoderm, *hb* is expressed in the serosa (orange) and as a cap in the posterior half of the embryo (blue) (T1) that eventually resolves into an expression band in the anterior (blue in T2 and T3). Later during the germband stage (T4), the anterior expression (blue) fades and a new *hb* expression arises in the posterior (purple). (**E**) Spatiotemporal dynamics of endogenous *hb* (green) and reporter gene *yellow* (red) expressions in hbA->*yellow Tribolium* embryos. Left panel: Time-staged embryos from 14 to 26 h AEL at 24 °C, in which mRNA transcripts (*hb*: green, *yellow:* red) were visualized using *in situ* HCR staining. Nuclear staining (Hoechst) is in gray. Right panel: Fluorescence signal along the dorsal-ventral axis was summed up to generate intensity distribution plots along the AP axis. (**F**) Detection of *de novo* transcription via *in situ* HCR staining of intronic *yellow*. Nuclear staining (Hoechst): blue; exonic *yellow (yellow* mRNA): red; intronic *yellow:* purple. Embryo outline shown in dashed line. (**G**) Live imaging snapshot of a hbA>MS2-*yellow*; aTub>MCP-mEmerlad *Tribolium* embryo. Diffuse mEmerald signal is observed in nuclei (outlined in white dashed line). mEmerald fluorescence is enriched at transcription sites (bright puncta: MS2-MCP signal). In all embryos shown: posterior to the right.

This system is capable of visualizing enhancer activity both in fixed and live embryos. To visualize aggregate enhancer activity in fixed embryos, *yellow* gene expression is visualized using *in situ* staining in embryos carrying the enhancer>MS2-*yellow* construct. To visualize *de novo* transcription in fixed embryos, an *in situ* probe against *yellow* intron is used instead. To visualize *de novo* transcriptional activity of an enhancer in live embryos, a male beetle carrying the enhancer>MS2-*yellow* construct is crossed with a female beetle carrying the aTub>MCP-mEmerald construct. If active, the enhancer should drive the expression of the MS2-*yellow* reporter in the progeny. The transcribed MS2 loops would then recruit aTub>MCP-mEmerald fusion proteins at the transcription site of the reporter gene, enriching the mEmerald fluorescent signal against the weak mEmerald background.

Via piggyBac transgenesis, we successfully generated a transgenic beetle line carrying the MCP-mEmerald construct, in which a ubiquitous mEmerald fluorescence is detected (**Figure 3C**). We then sought to test our enhancer>MS2-*yellow* reporter system, using a previously discovered *Tribolium* enhancer, hbA, that regulates the *Tribolium* gap gene *hunchback* (*hb*) (101). *hb* is expressed in multiple domains in the early *Tribolium* embryo: in the serosa, in an anterior domain, in a secondary posterior domain (shown in orange, blue, and purple, respectively in **Figure 3D**), and in the nervous system (not shown). Enhancer hbA drives the anterior expression of *Tribolium hb* (101) (blue in **Figure 3D**). Via piggyBac transgenesis, we successfully generated a transgenic beetle line carrying the hbA>MS2-*yellow* construct. Examining *yellow* expression using *in situ* hybridization chain reaction (HCR) (103) in early hbA>MS2-*yellow* embryos using both exonic (**Figure 3E,F**) and intronic (**Figure 3F**) probes, we confirmed that the *yellow* expression in hbA>MS2-*yellow* line is similar to the *mCherry* expression in a previously tested *hbA>mCherry* line (101).

To test the live imaging capability of our MS2-MCP system, we crossed the hbA>MS2-*yellow* and the aTub>MCP-mEmerald lines. Imaging early embryos of the progeny (hbA>MS2-*yellow*; aTub>MCP-mEmerald) (**Movie S1; Figure 3G**), we observed weak and diffuse mEmerald signal within the nuclei, and bright puncta at a rate of at most one puncta per nucleus. The bright mEmerald puncta are distributed along the AP axis initially as a cap that eventually refines into a stripe (**Supplementary Figure 9**), resembling the *yellow* expression of the hbA enhancer reporter visualized using *in situ* HCR staining (**Figure 3E,F**). We conclude, therefore, that such bright mEmerald puncta are mEmerald enrichments at transcribed MS2 loops, reflecting the *de novo* transcription driven by the hbA enhancer. Hence, both individual nuclei of the early *Tribolium* embryo and *de novo* transcription driven by the hbA enhancer can be visualized and detected in a single cross of hbA>MS2-*yellow* line and the MCP-mEmerald line, confirming our success in establishing an MS2-MCP enhancer reporter system that is capable of visualizing enhancer activity in live *Tribolium* embryos.

### Assessing the association between differential accessibility and enhancer activity

We then sought to use our enhancer reporter system to test putative enhancers suggested by our ATAC-seq analysis. In selecting a set of putative enhancers to test, we restricted our analysis to genomic regions around three genes, all involved in AP patterning in *Tribolium*: the gap genes *hb* (**Figure 4A**) and *Kr* (**Supplementary Figure 7A**) as well as the pair-rule gene *runt* (*run*) (**Figure 4B)**. Candidate enhancer regions were chosen based on the presence of accessible sites in any region along the AP axis and/or time point (**Materials and Methods**), whether or not they were differentially accessible.

**Figure 4.**
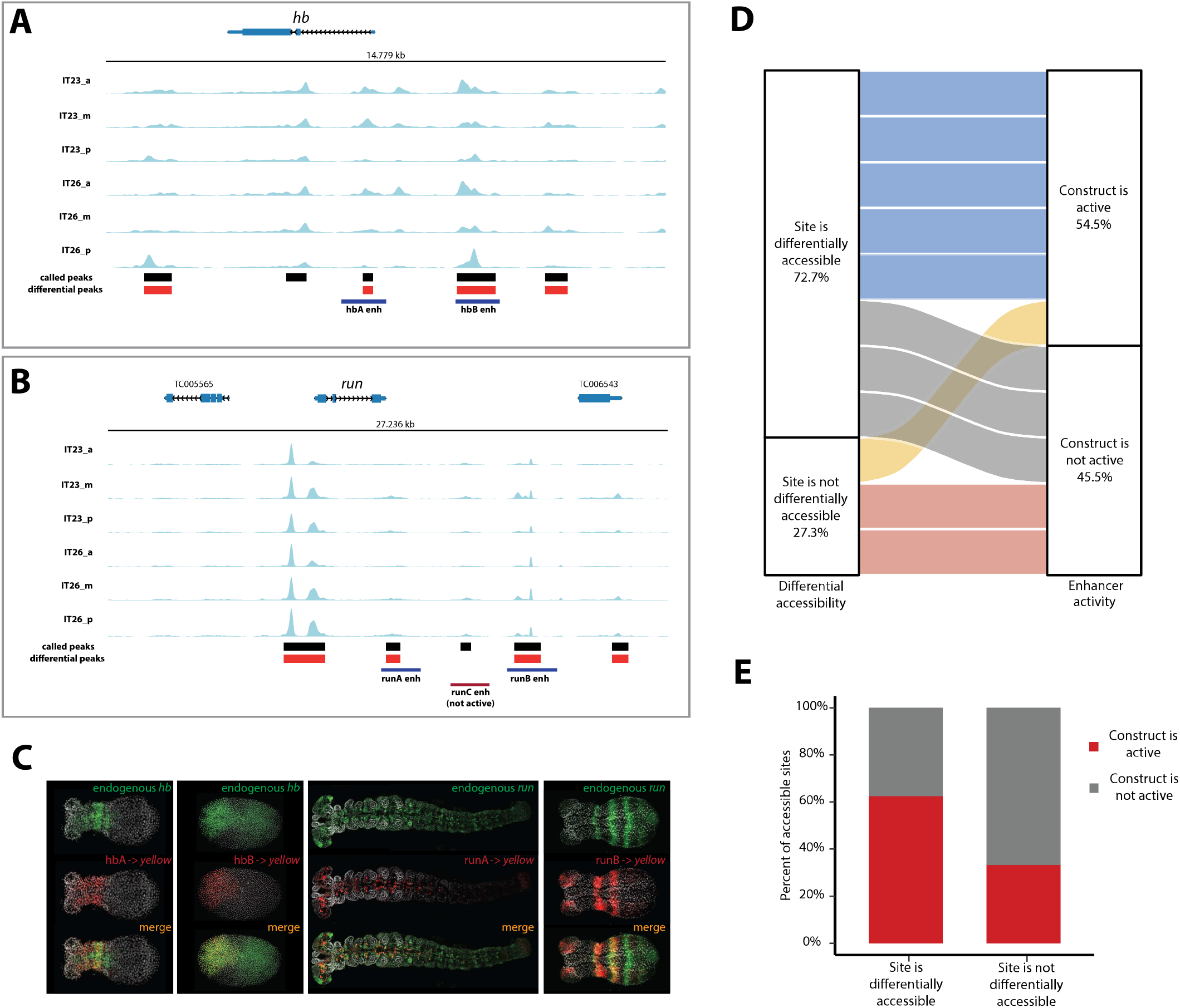
Correlation of enhancer activity with differential accessibility. (**A, B**) The ATAC profiles (two time points (IT23, IT26) with three embryo regions (a, m, p) per time point) for *hb* (A) and *run* (B). Tested enhancer regions at these loci are shown as boxes underneath the ATAC profiles. Differential accessible sites match well with active enhancer regions (purple boxes; red box: not active enhancer construct). ATAC tracks were created with pyGenomeTracks. (**C**) Enhancer reporter constructs for active enhancer regions shown in (A) and (B), in which mRNA transcripts were visualized using *in situ* HCR staining (endogenous gene expression: green, reporter gene expression: red, merge: yellow). Nuclear staining (Hoechst) is in gray. hbA drives reporter gene expression in a stripe (embryo in germband stage shown). hbB drives reporter gene expression in the serosa (embryo in blastoderm stage shown). runA drives reporter gene expression in a subset of the endogenous *run* CNS expression (embryo in late germband stage shown). runB drives reporter gene expression in stripes outside of the most posterior part of the embryo (embryo in germband stage shown). Posterior to the right. (**D**) The correlation between differential accessibility and construct activity was determined. Eleven enhancer constructs were analyzed: 54.5 % of constructs (six constructs) were active and 45.5 % of constructs were not active (five constructs). Five out of six active constructs are associated with sites that are differentially accessible, while one active construct overlaps with a site that is not differentially accessible. Two out of five not active constructs match sites that are not differentially accessible, while the remaining three not active constructs are associated with sites that are differentially accessible (see Supplementary Figure 7 for details). (**E**) Enhancer prediction efficiency of our enhancer prediction method based on differential peak analysis. Same enhancer constructs were analyzed as in (D). About 60 % of analyzed differential peaks were associated with active enhancer construct regions whereas in about 40 % of analyzed cases differential peaks could be found at not active enhancer construct regions. In contrast, about 70 % of analyzed non-differential peaks were associated with not active enhancer construct regions. About 30 % of analyzed non-differential peaks are associated with active enhancer construct regions.

Out of nine tested reporters, four successfully drove *yellow* expressions in the early *Tribolium* embryo (**Figure 4C**). While enhancer hbA drove an expression that overlaps with *hb* anterior expression, enhancer hbB drove an expression that overlaps with *hb* expression in the serosa (compare hbA and hbB activities in **Figure 4C**; see **Figure 3D** for the constituents of *hb* expression in *Tribolium*). Enhancer runA drove an expression that partially overlaps the endogenous *run* expression in the nervous system during the late germband stage (runA in **Figure 4C**). Enhancer runB drove a striped expression that overlaps endogenous *run* expression in the ectoderm, but neither the striped *run* expression in the mesoderm, nor the nervous system expression (runB in **Figure 4C**).

We then determined whether there is any association between differential accessibility and enhancer activity using our tested enhancer constructs, augmented with previously published *Tribolium* enhancers that regulate the genes *single-minded* (*sim*) and *short gastrulation* (*sog*) (104) (**Supplementary Figure 8**). In total, eleven enhancer constructs were analyzed, where 54.5 % of constructs (six constructs) were active and 45.5 % of constructs were not active (five constructs). We found that five out of six active constructs overlapped differentially accessible sites, while one active construct overlapped a site that was not differentially accessible (**Material and Methods**). Two out of five non-active constructs overlapped sites that were not differentially accessible, while the remaining three overlapped sites that were differentially accessible (**Figure 4D**; see **Supplementary Figure 7** for details). Therefore, about 60% of analyzed differentially accessible sites were associated with active enhancers whereas only 30% of analyzed constitutively accessible sites were associated with active enhancer regions (**Figure 4E**). These results support our hypothesis that differential accessibility is associated with enhancer activity.

### Testing the plausibility of the Enhancer Switching model

Next, we set out to test the plausibility of the Enhancer Switching model by examining the activity dynamics of some of the discovered enhancers. The model predicts that for a gene involved in generating gene expression waves, there exists two enhancers: (*i*) a ‘dynamic enhancer’ responsible for initiating the wave, and (*ii*) a ‘static enhancer’ responsible for arresting the wave into a stable gene expression domain(s) (**Figure 1C,D**).

Among the discovered enhancers in *Tribolium*, two enhancers are potentially involved in generating gene expression waves: hbA and runB. The expressions driven by both enhancers overlap with the expression waves of their corresponding genes: enhancer runB with the periodic waves of the pair-rule gene *run*, and enhancer hbA with the non-periodic wave of the gap gene *hb*. To test if the spatiotemporal dynamics driven by these enhancers conform with some of the predictions of the Enhancer Switching model, we first ran simulations of the model and used them to carefully analyze model predictions. Then, we used our enhancer reporter system to track the enhancer activity dynamics of runB and hbA in space and time using *in situ* HCR staining in carefully staged fixed embryos as well as using our MS2-MCP system in live *Tribolium* embryos. Finally, we compared our model predictions with the observed enhancer activity dynamics.

#### Careful examination of the predictions of the Enhancer Switching model

To carefully analyze the predictions of the Enhancer Switching model in space and time, we ran a simulation (**Movie S2**) of a 3-genes realization of the periodic version of the model (**Figure 1C**, where an oscillator is used as a dynamic module). Carefully analyzing model outputs for total activity of constituent genes, static enhancer reporters, and dynamic enhancer reporters revealed two characteristics of the spatiotemporal dynamics of their activities (**Figure 5A**). First, endogenous genes and reporters of their dynamic and static enhancers are all expressed in waves that propagate from posterior to anterior (**Figure 5A**). Second, expressions driven by dynamic enhancers progressively decrease in the posterior-to-anterior direction, matching the progressive decrease of the speed regulator concentration (**Figure 5A**). On the other hand, expressions driven by static enhancers progressively increase in the posterior-to-anterior direction, opposite to the direction of increase of the speed regulator concentration (**Figure 5A**). This is a natural consequence of the activating vs repressing effect of the speed regulator on dynamic vs static enhancers, respectively (**Figure 1C**).

**Figure 5.**
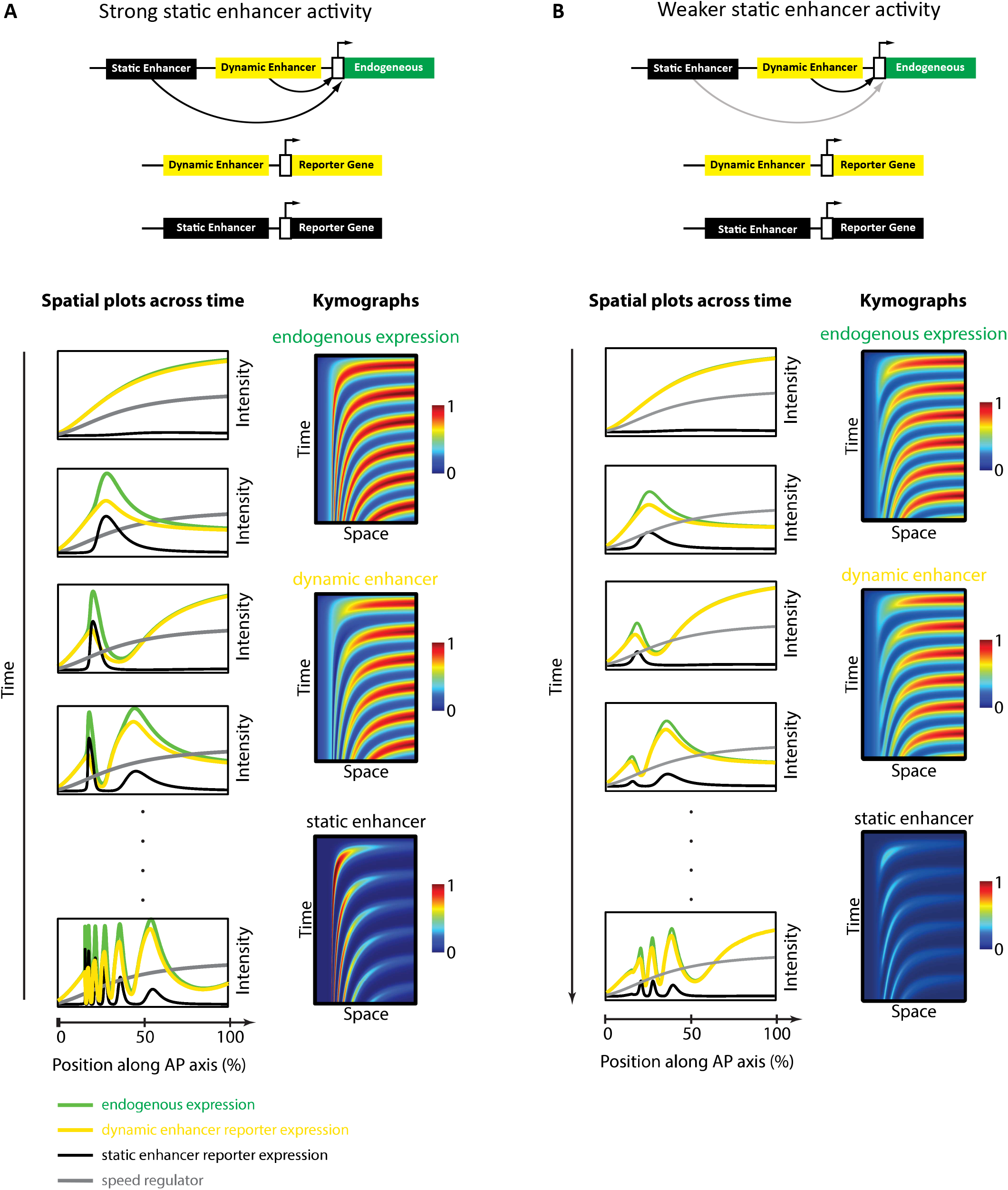
Simulation of the Enhancer Switching model with different static enhancer strengths. Shown simulation outputs of the Enhancer Switching model for a reporter gene driven by dynamic (yellow) or static (black) enhancers, as well as an endogenous gene driven by both dynamic and static enhancer (green). Two versions of the model were simulated and contrasted: with strong (A) vs weak (B) static enhancer activity. (**A**) Model simulation with strong static enhancer activity. Each wave of the endogenous gene expression follows first the dynamics of the dynamic enhancer and switches along space (in the tapering direction of the speed regulator, shown in gray) and time to the dynamics of the static enhancer to form a stable expression domain. (**B**) Model simulation with a weaker static enhancer: dynamic enhancer activity resembles endogenous gene expression pattern. Left panels: spatial plots across time. Right panels: Kymographs.

However, a minor complication arises when one considers a more realistic instantiation of the Enhancer Switching model. In our simulation of the model presented in **Figure 5A** (and in our simulations presented in previous publications (31,32,45)), we assumed that the stabilized gene expression domains at the anterior remain stable indefinitely (**Figure 5A**). However, we observe experimentally that such stable phase is transient, after which gene expression domains gradually fade (notice the progressive fading of the first *run* stripe after its stabilization at the anterior in **Figure 6A,B**). This effect can be implemented computationally by reducing the strength of the static enhancers (**Figure 5B**, **Movie S3**). In this case, the expression driven by a dynamic enhancer is very similar to the total expression of the gene expression wave: both arise maximally at the posterior and gradually fade as they propagate towards anterior (**Figure 5B**). On the other hand, the expression wave driven by the static enhancer remains unique, as it, for the most part, increases in the direction of its propagation (until it eventually fades; **Figure 5B**). This means that while it is easy to identify a static enhancer from its enhancer reporter activity, it is not as simple to identify a dynamic enhancer. In particular, an enhancer that drives an expression that arises maximally at the posterior and gradually fades as it propagates towards anterior might be either a dynamic enhancer or simply an enhancer that drives the entirety of the gene expression wave.

**Figure 6.**
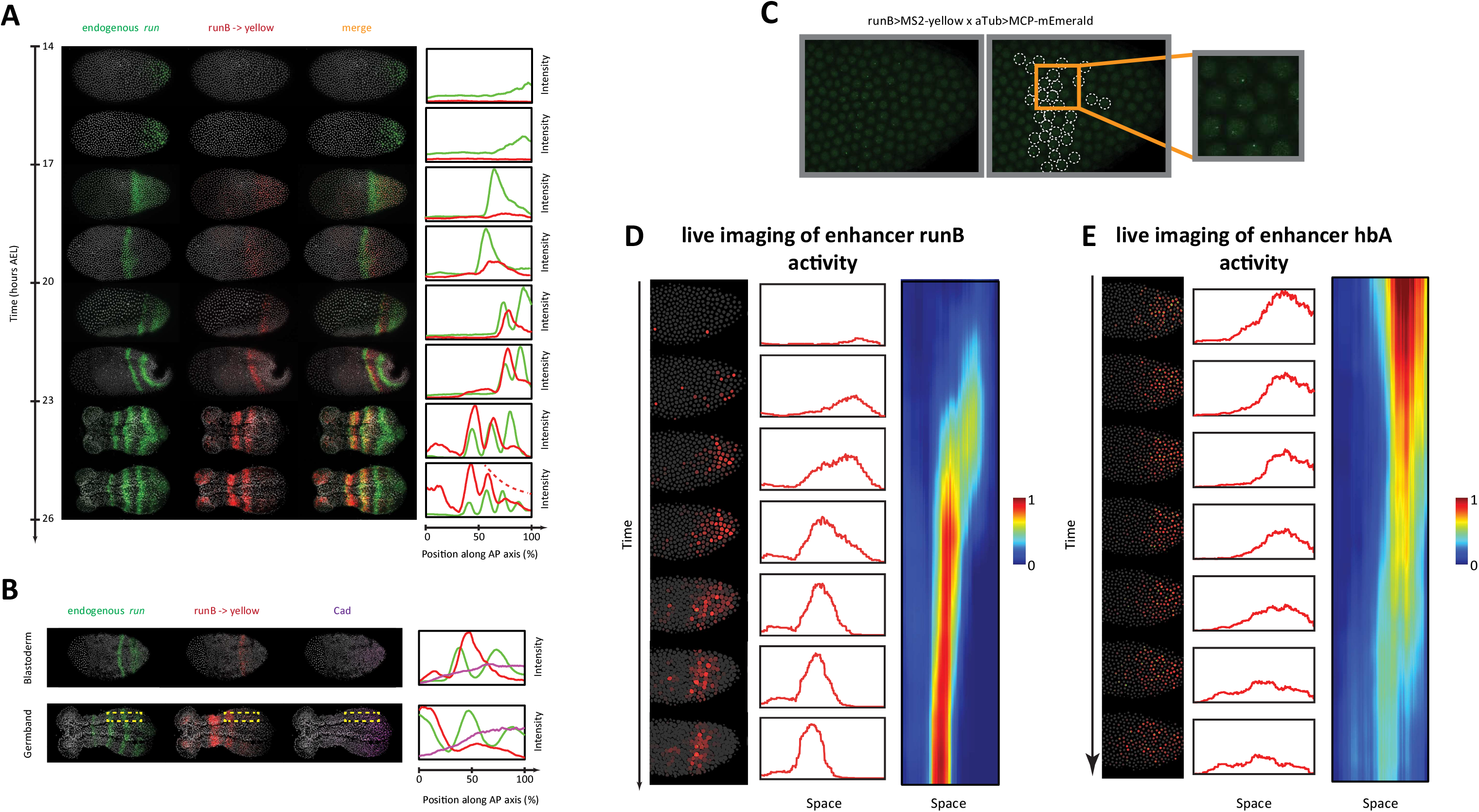
Analysis of enhancer activity dynamics using the MS2-MCP live imaging system. (**A**) Shown are spatiotemporal dynamics of endogenous *run* (green) and the reporter gene *yellow* (red) expression in runB->*yellow* embryos. Left panel: Time-staged embryos from 14 to 26 h AEL at 24 °C, in which mRNA transcripts (*run*: green, *yellow:* red) were visualized using *in situ* HCR staining. Nuclear staining (Hoechst) is in gray. Right panel: Intensity distribution plots along the AP axis. Both *run* and runB->*yellow* are expressed in waves that propagate from posterior to anterior. runB->*yellow* expression wave, however, starts weak posteriorly and progressively increases in strength as it propagates towards anterior, until it eventually overlaps with the stabilized *run* stripes anteriorly. (**B**) Left panel: *in situ* HCR staining for endogenous *run* (green) and reporter gene *yellow* expression (red) combined with antibody staining for Cad proteins (purple) in a runB->*yellow* blastoderm embryo (upper row) and a runB->*yellow* germband embryos (lower row). Nuclear staining (Hoechst) is in gray. Right panel: Intensity distribution plots along the AP axis for *yellow* expression in a whole *Tribolium* blastoderm (upper panel), and within the region indicated in dashed yellow in a *Tribolium* germband. Cad forms a posterior-to-anterior gradient in both balstoderm and germband embryos. runB activity increases progressively as Cad concentration drops towards anterior. (**C**) Live imaging snapshot of a runB>MS2-*yellow*; aTub>MCP-mEmerlad *Tribolium* embryo. Diffuse mEmerald signal is observed in nuclei. mEmerald fluorescence is enriched at transcription sites (bright puncta: MS2-MCP signal). Left panel: original snap shot; Middle: nuclei that exhibit MS2-MCP signal are outlined in white circles; Right: A close up to nuclei with MS2-MCP signal. (**D, E**) Tracking estimated mRNA activity driven by runB **(D)** and hbA **(E)**. Left panels (in both (D) and (E)): Snapshots across time from live embryo movies in which nuclei (shown in gray) are tracked and MS2 signals are averaged over time (using a moving average filter with a length of 10 movie frames) to estimate mRNA activity (shown in red). Middle panel: Intensity distribution plots along space for representative images in left panel. Right panel: A kymograph showing estimated mRNA activities of enhancer reporters across space and time. In all embryos shown: posterior to the right.

#### Examining the activity dynamics of enhancer runB using *in situ* HCR staining

To test the predictions of the Enhancer Switching model, we started by examining the spatiotemporal dynamics of one of the primary pair-rule genes, *run*, simultaneously with those of one of its enhancers that we discovered in this study, runB, using HCR in carefully staged embryos (**Figure 6A**). Endogenous *run* expression (green in **Figure 6A**) periodically arises from posterior and gradually propagates towards anterior, forming stable striped expression (that eventually fades). Concomitantly, runB>*yellow* is expressed as well in a periodic wave that propagates from posterior to anterior (red in **Figure 6A**). However, in contrast to endogenous *run* expression, the expression wave of runB>*yellow* appears weakly in the posterior and gradually strengthens as it propagates towards anterior. These observations are in line with the predicted enhancer dynamics of our model, in which runB acts as a static enhancer for *run*. First, runB drives gene expression waves that propagate from posterior to anterior. Second, runB activity progressively increases in the posterior-to-anterior direction, corresponding inversely with the progressive drop of concentration of the Caudal (Cad) gradient (**Figure 6B**), which has been suggested to act as a speed regulator for pair-rule and gap genes in *Tribolium* (30,31), and more generally, an evolutionary conserved posterior determinant in arthropods (along with Wnt) (64,105–108).

We notice, however, that the wave dynamics of runB>*yellow* expression are less discernible in posterior germbands (see **Figure 6A**, 23-26 hours AEL). We wondered if this is due to long degradation delays of *yellow* mRNAs. In line with this possibility, we noticed that mature runB>*yellow* stripes at anterior germbands are more stable and long-lived than those of endogenous *run* (**Supplementary Figure 10**). To circumvent this, we examined runB->*yellow* expression using an intronic probe against *yellow*, and indeed found that the intronic expression of runB->*yellow* is both discernible in the posterior germband and in line with endogenous *run* expression anteriorly (**Supplementary Figure 10**). This shows that, indeed, degradation delays of the reporter gene *yellow* is larger than that of endogenous *run*, leading to averaging out of *run* expression wave dynamics, a problem that can be alleviated using intronic *in situ* staining.

#### Examining the activity dynamics of enhancer runB using live imaging

To verify that runB indeed drives expression waves that propagate from posterior to anterior, we performed a live imaging analysis of runB activity using our MS2-MCP system in *Tribolium*. Crossing runB enhancer reporter line (runB>MS2-*yellow*) with our aTub>MCP-mEmerlad line, and imaging early embryos of the progeny, we observed bright mEmerald puncta distributed along the AP axis as a stripe (**Movie S4**; **Figure 6C**), an expression that resembles that of *yellow* in the same reporter line visualized using *in situ* HCR staining (compare **Figures 6A and 6C**). To characterize the spatiotemporal activity of runB enhancer, circumventing the ambiguity introduced by nuclear flow, we developed a computational strategy to: (*i*) track the nuclei of the early live *Tribolium* embryo, (*ii*) detect MS2 puncta, and (*iii*) associate the detected MS2 puncta to corresponding nuclei (**Materials and Methods**). Furthermore, to smoothen out the highly stochastic expression of *de novo* transcription, we applied a moving average window to MS2 signal across time to estimate an accumulated activity of the runB enhancer (**Movie S5; Materials and Methods**). Tracking the accumulated activity in space, after discounting nuclear flow, revealed that runB indeed drives a wave of activity that progressively increase in strength as it propagates from posterior to anterior (**Movie S6**; **Figure 6D**), fitting the role of a ‘static enhancer’ within our Enhancer Switching model.

#### Examining the activity dynamics of enhancer hbA using live imaging

Gap genes are expressed as well in waves in the *Tribolium* embryo, albeit in a non-periodic fashion. Gap gene waves are initialized by a pulse of *hb* expression that arises in the posterior of the blastoderm at 14 hours AEL, that eventually propagates towards anterior, clears from posterior, forming a stripe of *hb* expression at the anterior part of the embryo (**Figure 3D,E**).

Similar to our analysis of runB enhancer, we used our MS2-MCP system to estimate the accumulated mRNA signal driven by enhancer hbA, and tracked it in space and time in live *Tribolium* embryos (**Movie S7**, **Movie S8**). We found that enhancer hbA drives a wave of activity that propagates from posterior to anterior (**Figure 6E**). In contrast to enhancer runB, however, we found that enhancer hbA drives strong expression in the posterior that weakens as it propagates towards anterior (compare **Figures 6D** and **6E**). As concluded by our simulations of the Enhancer Switching model (**Figure 5B**), this indicates that hbA either drives the entirety of *hb* expression, or acts as a dynamic enhancer within the Enhancer Switching model.

## Discussion

In this paper, we established a framework for enhancer discovery in *Tribolium* using tissue- and time-specific ATAC-seq (**Figure 2**). We showed that differential accessible sites analysis across space and time yields a sizeable increase in enhancer prediction accuracy (**Figure 4**). We also developed an enhancer reporter system in *Tribolium* able to visualize dynamic transcriptional activities in both fixed and live embryos (**Figure 3**). Both our enhancer discovery and activity visualization systems are efforts to establish the AP patterning in *Tribolium* as a model system for studying dynamic gene expressions, especially gene expression waves, a phenomenon commonly observed during embryonic development (9,10,109). Although our experimental framework is suitable for exploring how enhancers mediate dynamic gene expression in an unbiased fashion, we set in this work to test the plausibility of a specific model: the ‘Enhancer Switching’ model (**Figure 1**), a scheme some of the authors have recently suggested (31,49) to elucidate how gene expression waves are generated at the molecular level. The model posits that each gene within a genetic clock or a genetic cascade is regulated by two enhancers: one ‘dynamic’ that induces rapid changes in gene activity, and another ‘static’ that stabilizes it. By modulating the balance between the potency of dynamic vs static enhancers, the tuning of the speed of gene regulation is achieved (**Figure 1C,D**). We first characterized the model’s predictions for the spatiotemporal activities of enhancer reporters of dynamic and static enhancers (**Figure 5**). The model predicts that reporter constructs of dynamic enhancers would drive a gene expression wave that progressively decreases in intensity in the direction of its propagation, whereas reporter constructs of static enhancers would drive a wave whose intensity increases in the direction of its propagation (**Figure 5A**). We then used our enhancer discovery framework to discover a number of enhancers regulating embryonic patterning in the early *Tribolium* embryo (**Figure 4**). One of these enhancers, runB, drove an expression pattern consistent with a role as a static enhancer for the pair-rule gene *run* (**Figure 6A–D**). Another enhancer, hbA, drove an expression pattern consistent with a role as a dynamic enhancer for the gap gene *hb* (**Figure 6E**). However, the expression pattern driven by hbA could be also interpreted as driving the entirety of *hb* expression (**Figure 6E**, **Figure 5B**). We present these findings as tentative support for the Enhancer Switching model, whereas a strong support requires: (1) finding enhancer pairs for several gap and pair-rule genes whose activity dynamics match those predicted for static and dynamic enhancers, and (2) verifying that the deletion of either dynamic or static enhancers result in phenotypes predicted by the model (**Figure 7**). Specifically, deleting a static enhancer should reduce the gene expression wave into an (almost) homogenously oscillating (or sequentially activating) domain at the posterior, that fails to resolve into gene expression bands anteriorly (**Figure 7B**), while deleting a dynamic enhancer should abolish the entire gene expression (**Figure 7C**). Future works should aim at testing these predictions, modifying the model, or finding alternative models.

**Figure 7.**
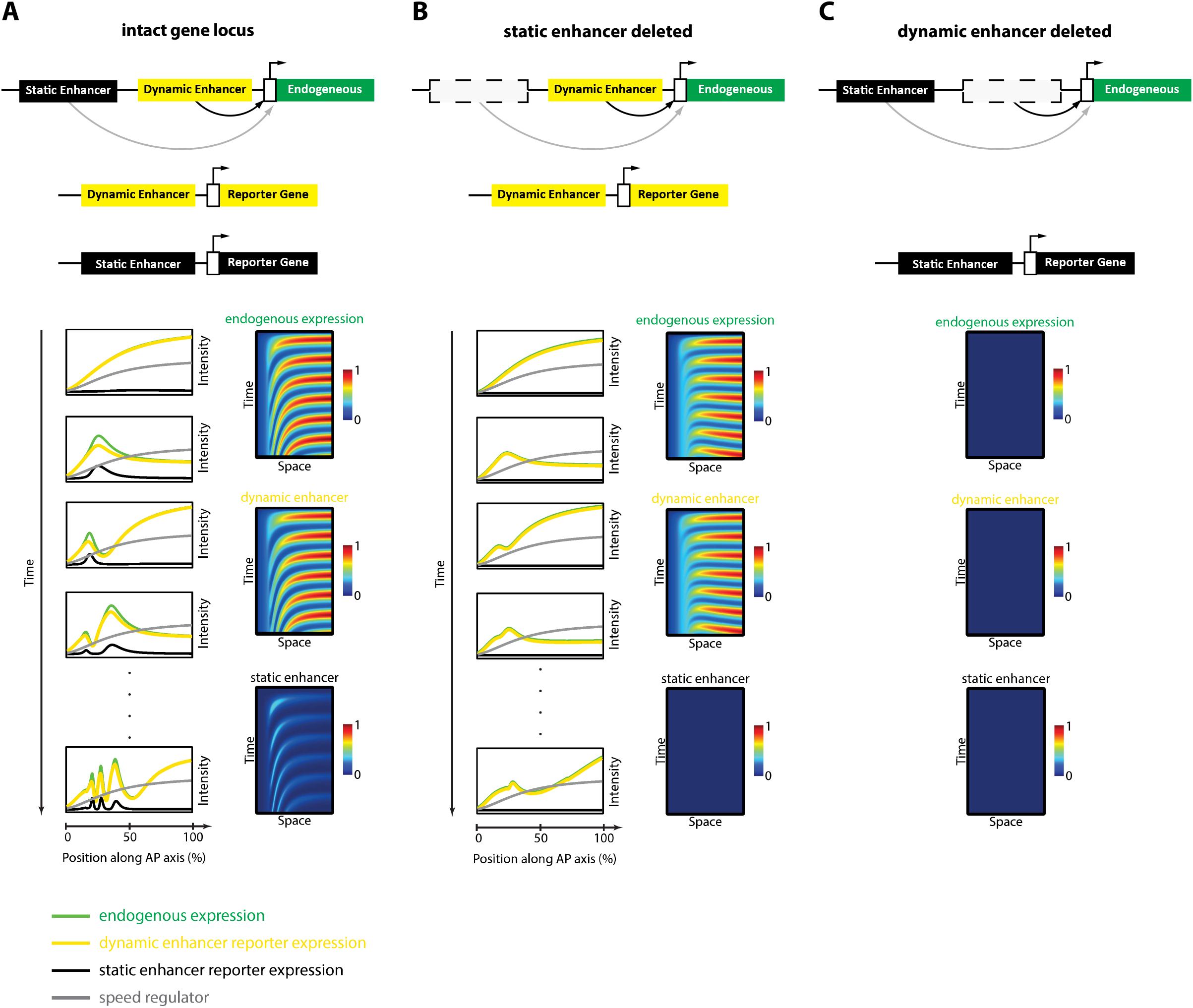
Simulation of the Enhancer Switching Model with deleted enhancers. Shown simulation outputs of the Enhancer Switching model under three experimental conditions: **(A)** an intact locus, **(B)** a locus in which the static enhancer is deleted, and **(C)** a locus in which the dynamic enhancer is deleted.

The long-term goal of establishing the experimental framework presented in this paper is to understand the regulation of dynamic gene expressions at the molecular level. The model system we adopted exhibits, however, very particular kind of dynamics: wave dynamics. This sounds like attempting to solve a fringe problem, since the most notable and well-known instance of this problem has been documented during vertebrate somitogenesis, in which case it is unclear if such wave dynamics are of any functional significance. Indeed, the predominant model of somitogenesis, namely the ‘Clock-and-Wavefront’, can do without such wave dynamics. Furthermore, gene expression waves observed during somitogenesis are periodic, making the problem even more of a fringe case, as there are far less cases of periodic gene expression patterns in development than non-periodic ones. However, upon reviewing experimental observations in other model systems, one finds strong indications that such wave dynamics, especially in their non-periodic version, are more prevalent than has previously thought, albeit described in varied terminologies. For example, Hox genes, which are expressed in a non-periodic pattern along the AP axis of vertebrates, have been described to ‘spread’ anteriorly upon their emergence from the posterior (18). Similarly, neural fate-specifying genes have been described to arise in a variable temporal order along the dorsoventral axis of the vertebrate neural tube, and so are effectively expressed as a non-periodic wave (22). Furthermore, the functional importance of such wave dynamics is manifest in embryonic tissues that undergo no or limited elongation during the patterning process, where a pure Clock-and-Wavefront model fails to explain the patterning process (9) (e.g. the vertebrate neural tube and limb bud, the vertebrate embryo at the early phase of somitogenesis (110), and the blastoderm of short-germ insects). Even for elongating tissues that can be patterned by a classical Clock-and-Wavefront mechanism, wave dynamics have been suggested to foster robustness for the patterning process (30,111,112).

As we set our experimental framework as means to study gene expression waves in development, we should note that we have only discussed one type of waves, namely ‘kinematic waves’ (also called ‘phase waves’ or ‘pseudo-waves’) (10,113). A kinematic wave propagates without the need of ‘diffusion’ or cell-to-cell communication, and gives the appearance of spreading across the tissue due to differences in gene expression timing between different cells. This type of wave can mediated by the Speed Regulation model discussed in this study (**Figure 1A, and 1B**). A wave, however, could be also a ‘trigger’ wave, which propagates with the help of cell-to-cell communication (10). One key experiment to differentiate between a trigger versus a kinematic wave is to insert a barrier along its path. The barrier should block a trigger wave, while a kinematic wave would appear to continue propagating across the barrier. A barrier experiment of that sort indicated that the periodic gene expression waves observed during somitogenesis are indeed of the kinematic type (11). Another key feature that differentiates a trigger from a kinematic wave is the spatial variability of the wave’s wavelength. Kinematic waves of the sort induced by the Speed Regulation model has a key signature: it emanates as wide bands, that progressively shrink in the direction of its propagation, and hence the wave’s wavelength is variable across space (**Figure 1A,B**). A trigger wave, on the other hand, is typically of a fixed wavelength. Most of the waves described during embryonic development (including gap and pair-rule waves in *Tribolium*) shrink in the direction of their propagation, and thus exhibit the appearance of kinematic waves (see for example **Figure 6D,E**).

Once a gene expression wave is determined to be kinematic, mechanistically explaining how it is produced reduces to elucidating how the speed of gene regulation can be tuned by a morphogen gradient (according to Speed Regulation model: **Figure 1A,B**). Whereas the phenomenon of gene expression waves is of significance within the domain of embryonic pattern formation, the problem of how to tune the speed of gene regulation has a wider applicability. For example, the temporal progression of different developmental stages is species-dependent and, hence, needs to be tunable by evolution (114), the timing of developmental progression in insects is controllable by the steroid hormone ecdysone (115), and the time needed for progenitor cells to differentiate needs to be tuned to control (relative) population/organ sizes, which furthermore needs to be adjustable by evolution (116–119). Thus, the model problem we are introducing in this paper has wide applicability within and beyond the domain of embryonic pattern formation.

In this paper, we considered the Enhancer Switching model as a molecular realization for how to modulate the timing of gene regulation. The central idea behind the model is that tuning the timing of gene regulation is the result of setting the balance between two GRNs (**Figure 1C**): one GRN mediates rapid changes in gene regulation (termed ‘dynamic GRN’), and another stabilizes gene expressions (termed ‘static GRN’). This scheme could be realized by just one enhancer per gene, if such an enhancer is able to switch its wiring depending on the concentration of a graded molecular factor (the speed regulator). The phenomenon of one enhancer exerting multiple functions has indeed been observed and termed ‘enhancer pleiotropy’ (120–124). However, we suggested a molecular strategy that uses a separate enhancer for each wiring scheme (**Figure 1D**) (31). This strategy, we believe, is more molecularly feasible and ensures modularity and evolvability (31). Indeed, the complex regulation of many genes in development are usually undertaken by several enhancers, each encoding simple regulatory logic (125–127).

While the evidence for the Enhancer Switching model is still sketchy, parallels (or evolutionary vestiges) to it is evident in *Drosophila*. After an initialization phase, the striped expression of *Drosophila* pair-rule genes stabilizes and undergoes frequency doubling. The switching between the initialization to stabilization mode of pair-rule regulation is mediated by the switch of early to late acting enhancers. Interesting, this switch seems to be mediated by timing factors encoded by two pioneer factors: Zelda and Opa, where Zelda is responsible for activating the early network and Opa for the late network (128–130). Indeed, in *opa* mutants, the frequency doubling of pair-rule genes is lost in *Drosophila* (128). Interestingly, *cad* (potentially along with *zelda*), and *opa* were found to be activated sequentially in the *Drosophila* embryo (and *Nasonia’s* (131)), reflecting a similar sequential activation in space and time in the *Tribolium* embryo (47,132). This gives rise to a possible unified model for early-to-late enhancer switching in insects. In this model, early/posterior expression of AP patterning genes are mediated by early/dynamic enhancers, while late/anterior expressions are mediated by late/static enhancers. This transition is mediated ancestrally by Wnt/Cad gradient, possibly indirectly through Zelda and Opa, where Zelda is responsibly for activating early/posterior/dynamic enhancers, whereas Opa is responsible for activating late/anterior/static enhancers (9). Such late enhancers might, however, play a mere maintaining role as indeed commonly believed, while a static enhancer as suggested by the Enhancer Switching model has a key functional role in sculpting the waves. Such functional importance can only be elucidated using an enhancer deletion experiment (**Figure 7**).

Mechanisms for tuning the speed of gene regulation other than the Enhancer Switching model, however, are also possible. For example, histone modifications have been shown to influence the timing of gene activation or repression (133). Although such a mechanism does not offer complete tunability of the speed of gene regulation (since histone modifications usually specifically influence either gene activation or repression), it can explain certain phenomena; for example, controlling the timing of sequential activation (but not inactivation) of fate specifying genes (116,117,133). Modulating enhancer accessibility could be yet another mechanism for tuning the timing of gene regulation (115). However, such a scheme would require a yet unknown mechanism for stably maintaining gene expression while enhancer accessibility is reduced, without the help of a stabilizing enhancer (otherwise, it would be simply an instance of the Enhancer Switching model). Such possibilities could also be explored using our model system, if complemented with genomic assays for histone modifications.

In yet another hypothesis for how to generate gene expression waves, the timescale of the genetic process (be it a genetic clock or a cascade) is tuned by modulating protein production time delay or the coupling strength between cells (134–140). However, in *Tribolium*, reactivating the gap gene *hb* all over the embryo resulted in two distinct responses (32): reactivation of the genetic gap cascade in the posterior (where the speed regulator Wnt/Cad has high concentration), whereas the expression of already formed gap gene domains at the anterior is erased. This indicates that the switch in behavior of gap gene regulation between the anterior vs posterior in the early *Tribolium* embryo is not due to a simple change either in protein and transcript degradation rates or cell-to-cell coupling strengths, but due to a difference in the genetic makeup between these two regions, giving more credit to our Enhancer Switching model.

## Data Availability

ATAC-seq tracks were included in the iBeetleBase Genome Browser (https://ibeetle-base.uni-goettingen.de/genomebrowser/) (141).

## Acknowledgement

This work is supported by a DFG grant (EL 870/2-1) to EE, and a doctoral fellowship from the German Academic Scholarship Foundation to CM. We thank Rodrigo Nunes da Fonseca, Emilila Esposito, Alexander Aulehla, and Shelby Blythe for providing valuable tips for ATAC-seq library preparation.

## Materials and Methods

### Beetle Cultures

Beetle cultures were reared on flour supplemented with 5 % dried yeast in a temperature- and humidity-controlled room at 24 °C. To speed up development, beetles were reared at 32 °C.

### PiggyBac Enhancer Reporter Constructs

A piggyBac plasmid with the 3 x P3-mCherry/mOrange marker construct and multiple cloning sites (77) was used to generate all enhancer constructs in this study. For enhancer constructs, putative enhancer regions, the *Drosophila* synthetic core promoter (101,142), and the MS2-*yellow* reporter gene (97,99) were amplified by PCR, assembled through ligation, and inserted into the multiple cloning site of the piggyBac plasmid. Used primers are listed in **Table S1**.

### Creation of the MCP-mEmerald construct

An artificial sequence, consisting of (i) an AscI restriction enzyme site, (ii) the *Tribolium castaneum* Georgia2 (GA2) background strain-derived (143) tubulin alpha 1-like protein (aTub) promoter (144), (iii) a *Tribolium castaneum* codon-optimized open-reading frame consisting of (a) the SV40-derived nuclear localization signal (NLS) tag (145) coding sequence, (b) the human influenza hemagglutinin (HA) tag (146) coding sequence and (c) the bacteriophage MS2 coat protein (MCP) (147) coding sequence, and (iv) a NotI restriction enzyme site, was *de novo* synthesized and inserted into the PacI/SacI restriction enzyme site pair of pMK-T (Thermo Fisher Scientific) by the manufacturer. The resulting vector was termed pGS[aTub’NLS-HA-MCP]. The sequence was excised with AscI/NotI and inserted into the backbone of the accordingly digested pACOS{#P’#O(LA)-mEmerald} vector (77). The resulting vector was termed pAGOC{aTub’NLS-HA-MCP-mEmerald}, contained (i) an expression cassette for mEmerald-labeled (148) and NLS/HA-tagged MCP, (ii) the piggyBac 3’ and 5’ inverted terminal repeats (149), as well as (iii) mOrange-based (150) and mCherry-based (151) eye-specific (152) transformation markers, and was co-injected with the standard piggyBac helper plasmid (153) into *Tribolium castaneum* embryos following standard protocols (154,155) to achieve germline transformation.

### *Tribolium* Transgenesis

PiggyBac constructs were transformed into vermilion^white^ (156) with mCherry/mOrange as visible makers. Germline transformation was carried out using the piggyBac transposon system (153,157).

### Egg Collections for Developmental Time Windows

Developmental time windows of three hours were generated by incubating three hours egg collections at 24 °C for the desired length of time. Beetles were reared in flour supplemented with 5% dried yeast.

### *In Situ* Hybridization and Imaging of Fixed Embryos

*In situ* hybridization was performed using the third generation *in situ* hybridization chain reaction (HCR) method (103). All probe sets and hairpins were ordered at Molecular Instruments. Lot numbers of probe sets are as follows: PRA978 (*run* mRNA), PRA979 (*hb* mRNA), PRC655 *(yellow* mRNA), and PRE723 *(yellow* intron). Images were taken with a Leica SP5 II confocal. A magnification of 20x or 63x was used at a resolution of 2,048 x 1,024. Images were processed and enhanced for brightness and contrast using Fiji (158).

### Live Imaging

aTub>MCP-mEmerlad females were crossed with hbA>MS2-*yellow* or runB>MS2-*yellow* males. Three hours egg collections were generated and incubated for eleven (crossing with hbA) or fourteen hours (crossing with runB) at 24 °C. Embryos were dechorionated by immersion in 1 % bleach for 30 s twice. Embryos were mounted using the hanging drop method and covered with halocarbon oil 700 (Sigma). Time-lapse movies were taken by capturing 41 planes every 3 min over ~ 6 h at 21 °C with a Leica SP5 II confocal. A magnification of 63X was used at a resolution of 1,024 x 900. To produce unprocessed live imaging movies (**Movies S1 and S4**), a maximum Z-projection is applied to the image sequence in Fiji.

### Computational Processing and Analysis of Live Imaging Movies

To characterize the transcription dynamics driven by enhancer MS2 enhancer reporters in live embryos, circumventing the ambiguity introduced by nuclear flow, we developed a computational strategy to: (1) segment nuclei, (2) detect MS2 spots and estimate their intensity, (3) associate MS2 spots to nuclei and track nuclei over time, and estimate mRNA intensity.

#### (1) Segmenting nuclei

In Fiji, stacks were first maximum intensity projected. Contrast was enhanced using the CLAHE plugin with a block size of 128. Nuclei were then detected as local maxima, disregarding maxima with an intensity below half the image maximum intensity. Detected maxima were used as seed points for the watershed algorithm to retrieve nuclei outlines.

#### (2) MS2 spot detection

In Fiji, MS2 spots were detected as local 3D maxima after applying a 3D Difference-of-Gaussians filter. Its parameters, the standard deviations of the Gaussians, tolerance (the minimum intensity difference between neighbor spots, analogous to the ImageJ ‘Find Maxima’ implementation) and a lower threshold (to disregard spots with low intensity) were set empirically.

#### (3) Tracking nuclei and MS2 spots over time and mRNA estimation

We used strategies similar to those described in (41) using Matlab.

### ATAC-seq Library Preparation

Embryos of the nGFP line (29) were collected in flour supplemented with 5% dried yeast for 3h and incubated for desired time length at 24 °C. Embryos were dechorionated by immersion in 1 % bleach for 30 s twice. Selected embryos were dissected into three parts (anterior, middle, and posterior). For each biological replicate, three of the same embryo parts were pooled, and three replicates were prepared for each sample condition. Library preparation was performed as previously described (159,160). Tagmentation was performed for 8 min. ATAC-seq libraries were sequenced on an Illumina NovaSeq 6000 at the Novogene Cambridge Genomic Sequencing Centre. 2×150 bp paired-end Illumina reads were obtained for all sequenced ATAC-seq libraries.

### ATAC-seq Data Pre-processing

Sequencing reads were trimmed with cutadapt (161) with parameters “-u 15 -U 15 -q 30 -m 35 --max-n 0 -e 0.10 -a CTGTCTCTTATA -A CTGTCTCTTATA” to remove adapter sequences and mapped to the *Tribolium castaneum* reference genome (Tcas5.2, GCA_000002335.3) with BWA-MEM (version 0.7.12-r1039 (162). Next, read duplicates were marked with Picard MarkDuplicates (version 2.15.0, Picard Toolkit. 2019. Broad Institute, GitHub Repository. https://broadinstitute.github.io/picard/; Broad Institute.). Low quality and duplicated reads were filtered using samtools view (version 1.10, (163)) with parameters “-F 1804 -f 2 -q 30”. To flag regions that appear to be artifacts, we generated a blacklist using a strategy similar to the one developed by the ENCODE Project (164). Specifically, the genome was first divided into non-overlapping 50bp bins. Next, the BAM files containing the filtered mapped reads were converted into BigWig files using BAMscale (version 1.0, (165)) with parameters “scale --operation unscaled --binsize 20 --frag”. Using the resulting BigWig files, the mean signal for each bin was computed across all sequencing libraries. Finally, bins with a mean signal equal to or greater than 100 were flagged as artifacts and included in a “blacklist”. The threshold was determined by visual inspection of the distribution of the mean signals. Reads mapping to genomic regions in the blacklist were filtered out using samtools view (version 1.7, (163)) with parameters “-L” and “-U”.

Peaks were called on individual replicates using macs3 (v3.0.0a7, https://github.com/taoliu/MACS; (166)) callpeak with parameters “-g 152413475 -q 0.01 -f BAMPE”. The sets of peaks called in each sample were then compared to each other and merged if they overlapped by at least 1 bp. Only merged peaks supported by peaks called in at least two different samples and on scaffolds assigned to linkage groups were considered for subsequent analyses. These peaks are further referred to as the set of “all” (consensus) chromatin accessible sites.

Chromatin accessible sites were annotated with HOMER (version 4.11.1, (167)) using the annotatePeaks.pl function.

### Genomic tracks

Normalized ATAC-seq coverage tracks were generated with BAMscale (version 1.0, (165)) using parameters “scale --frag --binsize 20 --smoothen 2”. The tracks of different biological replicates of the same sample were then merged with the UCSC tools bigWigMerge and bedGraphToBigWig (http://hgdownload.cse.ucsc.edu/admin/exe/linux.x86_64/), and visualized with pyGenomeTracks (version 3.7, (168,169)).

### Differential Accessibility Analysis

Differential accessibility analysis of the sites between different regions of the germband and/or time points was performed using the edgeR (version 3.36.0, (170)) and DESeq2 (version 1.34.0, (171)) methods within the DiffBind (version 3.4.11, (172), http://bioconductor.org/packages/release/bioc/vignettes/DiffBind/inst/doc/DiffBind.pdf). Peaks with a false discovery rate (FDR) less than or equal to 0.05 for edgeR and/or DESeq were considered significant. Read counts for each peak were quantified with the dba.count() function, with parameters “score=DBA_SCORE_TMM_MINUS_FULL, fragmentSize=171, bRemoveDuplicates=TRUE”. Briefly, this normalizes the read counts using Trimmed Mean of M-values (TMM, (173)) scaled by the full library size. We further refer to these values as “accessibility scores”. Sites that had a mean accessibility score larger than the median and a standard deviation larger than the 3rd quartile were defined as the “most variable accessible sites”.

#### Clustering

The z-score normalized accessibility scores of the differentially accessible sites were clustered using k-means (with k=6) as implemented in the kmeans() R function. The function was executed with default parameters except for “nstart=1000”, which initializes the algorithm with 1000 different sets of random centers and settles for those giving the best fit.

Complete linkage on the pairwise Pearson correlation distances (1-Pearson correlation coefficient) was performed with the hclust() R function to hierarchically cluster the differentially accessible sites based on their average accessibility scores across the replicates of each sample group. The pheatmap (version 1.0.12, Raivo Kolde (2019). pheatmap: Pretty Heatmaps. R package version 1.0.12. https://CRAN.R-project.org/package=pheatmap) R package was used for visualization using row-scaling.

#### Functional analysis

Functional enrichment analysis was carried out with the PANTHER API. Specifically, the “statistical enrichment test” tool was used with the “GO Biological Process complete” *Tribolium castaneum* database. Overrepresented GO terms/pathways were determined using default parameters (Fisher’s exact test, False Discovery Rate (FDR) < 0.05). All the positive overrepresented GO biological process (BP) terms were considered. The API was last accessed on August, 2022.

#### Motif analysis

Motif enrichment analysis was carried out using XSTREME with default parameters except for “--streme-totallength 4000000 --meme-searchsize 100000 --ctrim 500 --streme-nmotifs 5” (version 5.4.0, (174)). Motifs discovered *de novo* were compared to the motifs in the insects JASPAR 2022 CORE non-redundant database (175) to identify putative transcriptional regulators. Sequences in which soft-masked nucleotides comprised 95% or more were not considered for the analysis.

#### Overlap between Differentially Accessible Sites and Constructs

A construct was considered to be overlapping with a site if at least 90% of the site overlapped with the construct.

### Computational Modeling

For the Enhancer Switching Models, we used computational algorithms described in (31,32,49), using the Matlab programs provided in these publications with minor modifications.

---

**Movie S1. MS2 live imaging of hbA enhancer reporter.** Live imaging of a “hbA>MS2-*yellow*; aTub>MCP-mEmerlad” *Tribolium* embryo during the blastoderm stage. NLS signal within the aTub>MCP-mEmerald construct mediates a weak and diffuse mEmerald signal within nuclei. Upon transcription, MS2 loops within the hbA>MS2-*yellow* construct recruit MCP-mEmerald fusion proteins at transcription sites, resulting in mEmerald bright puncta. Here bright mEmerald puncta are observed throughout the posterior end of the blastoderm, reflecting transcriptional activity of enhancer hbA in the early *Tribolium* embryo. Posterior to the right.

**Movie S2. Simulation of the Enhancer Switching Model with strong static enhancer activity**. Shown are outputs of a computer simulation of the Enhancer Switching model with strong static enhancer activity. Activity dynamics of reporter genes driven by the dynamic and static enhancers are shown in yellow and black, respectively. Activity dynamics of endogenous gene expression driven by both the dynamic and static enhancers are shown in green. Speed regulator gradient is shown in grey.

**Movie S3. Simulation of the Enhancer Switching Model with weak static enhancer activity.** Shown are outputs of a computer simulation of the Enhancer Switching model with wea static enhancer activity. Activity dynamics of reporter genes driven by the dynamic and static enhancers are shown in yellow and black, respectively. Activity dynamics of endogenous gene expression driven by both the dynamic and static enhancers are shown in green. Speed regulator gradient is shown in grey.

**Movie S4. MS2 live imaging of runB enhancer reporter**. Live imaging of a “runB>MS2-*yellow*; aTub>MCP-mEmerlad” *Tribolium* embryo during the blastoderm stage. NLS signal within the aTub>MCP-mEmerald construct mediates a weak and diffuse mEmerald signal within nuclei. Upon transcription, MS2 loops within the runB>MS2-*yellow* construct recruit MCP-mEmerald fusion proteins at transcription sites, resulting in mEmerald bright puncta. Here bright mEmerald puncta are observed initially to be distributed as a posterior cap that eventually propagates towards anterior to form a stable band. Posterior to the right.

**Movie S5. Estimated mRNA transcription driven by enhancer runB in the early *Tribolium* embryo.** Shown is a live imaging movie of a “runB>MS2-*yellow*; aTub>MCP-mEmerlad” embryo (same as in Movie S4) computationally processed to show an estimation of accumulated mRNA abundance driven by enhancer runB (red) as well as MS2-mEmerald signal (reflecting *de novo* transcription; green). Posterior to the right.

**Movie S6. Plots of estimated mRNA transcription dynamics driven by enhancer runB across space and time.** Shown is a dorsoventral projection of a tracked spatiotemporal activity of enhancer runB (same embryo as in **Movies S4,S5**). Horizontal axis: AP axis; posterior to the right.

**Movie S7. Estimated mRNA transcription driven by enhancer hbA in the early *Tribolium* embryo.** Shown is a live imaging movie of a “hbA>MS2-*yellow*; aTub>MCP-mEmerlad” embryo (same as in Movie S1) computationally processed to show an estimation of accumulated mRNA abundance driven by enhancer hbA (red) as well as MS2-mEmerald signal (reflecting *de novo* transcription; green). Posterior to the right.

**Movie S8. Plots of estimated mRNA transcription dynamics driven by enhancer hbA across space and time.** Shown is a dorsoventral projection of a tracked spatiotemporal activity of enhancer hbA (same embryo as in **Movies S1,S7**). Horizontal axis: AP axis; posterior to the right.

**Supplementary Figure 1. Correlation between ATAC-seq sequencing libraries**. Mapped sequencing reads of each biological replicate were used to calculate correlation between pairs of replicates. The values represent the Pearson correlation coefficients between pairs of sequencing libraries.

**Supplementary Figure 2. Upset plot comparing the number of sites identified in samples corresponding to IT23 and IT26 as well as along the AP axis**. (**A**) Plot corresponding to IT23 and IT26. (**B**) Plot corresponding to samples along the AP axis (a, m, and p). The number of sites in each of the sets considered is represented by the width of the bars in the bar chart at the bottom left. Each bar in the bar chart at the top represents the number of sites in an intersection. The intersection is indicated by filled circles below each bar. Note that the intersection sets are disjoint.

**Supplementary Figure 3. Annotation of consensus sites**. A total of 12069 consensus sites were analyzed. Consensus sites overlap with promoter-TSS (22.3 %), TSS (8.5 %), exons (23.5 %), introns (30.2 %), and intergenic regions (15.5 %), respectively. TSS: Transcription Start Site.

**Supplementary Figure 4. Analysis of differentially accessible sites**. (**A**) The accessibility of sites was compared between three different parts of the germband (anterior, middle, and posterior) at two time points (IT23 and IT26). All possible 15 comparisons were considered. A total of 3106 sites were differential accessible. (**B**) For the same part of the embryo, 132 sites were differentially accessible between two time points. (**C**) A total of 2109 sites were differentially accessible between two different parts of the germband at a given time point.

**Supplementary Figure 5. Motif Analysis.** Enrichment of TF motifs in accessible sites depending on their accessibility (y-axis, see Figure 1D for details). The size of the dots represent the fraction of sequences with one or more motif occurrences. The TFs (if known) are indicated on the x-axis.

**Supplementary Figure 6. Functional enrichment analysis of the clusters of differentially accessible sites**. Clustering of all differentially accessible sites across the AP axis and time points. All clusters are associated with “developmental process” and “anatomical structure development”, while clusters 1 and 3 are specifically related to “pattern specification process” and “regionalization” and cluster 3 is specifically associated with “anterior/posterior pattern specification”.

**Supplementary Figure 7. Genomic tracks of analyzed enhancer reporter constructs.** Genomic tracks of analyzed enhancer reporter constructs. ATAC profiles (two time points (IT23, IT26) with three embryo regions (a, m, p) per time point) are shown for (**A**) *Kruppel* (*Kr*), (**B**) *short gastrulation* (*sog*), and (**C**) *single-minded* (*sim*). Analyzed enhancer regions at these loci are shown as purple (active enhancer region) or red (not active enhancer region) boxes underneath the ATAC profiles. KrA enh - KrD enh were tested in *Tribolium* in this work. Other enhancer regions were evaluated in a cross-species context in *Drosophila* (104). Differential accessible sites match well with active enhancer region of *sim* (C). No differential accessible site overlap with the previously described active enhancer region of *sog* (B). Not active KrB enh region is not overlapping with differential accessible sites, while overlaps are observed between differential accessible sites and not active KrA enh, KrC enh, and KrD enh regions (A). ATAC tracks were created with pyGenomeTracks.

**Supplementary Figure 8. Simulations with different static enhancer strength.** Computational simulation of reporter gene expression driven by dynamic (yellow) or static enhancer (black), respectively as well as endogenous gene expression (green) driven by both, dynamic and static enhancer. Analyzed was the spatiotemporal pattern of gene expression. (**A**) No static enhancer: dynamic enhancer strength d=3, static enhancer strength c=0 (same as in Figure 7B, deleted static enhancer). Each wave of the endogenous gene expression follows the dynamics of the dynamic enhancer. Due to the lack of static enhancer activity, endogenous gene expression waves fail to stabilize as stable expression domains. (**B**) Very weak static enhancer: d=3, c=0.5. Each wave of the endogenous gene expression follows the dynamics of the dynamic enhancer. Due to very weaker static enhancer activity, dynamic enhancer activity resembles whole gene expression pattern (compare to Figure 5B). Endogenous gene expression waves are impaired in forming stable expression domains. (**C**) Weak static enhancer: d=3; c=1. Dynamics of the endogenous gene expression are mainly governed by dynamic enhancer activity. Weak static enhancer activity stabilizes endogenous gene expression waves as stable expression domains. (**D**) Intermediate strong static enhancer: d=3, c=1.5. Each wave of the endogenous gene expression follows first the dynamics of the dynamic enhancer and switches along space to the dynamics of the intermediate strong static enhancer to form a stable expression domain. Left panel: Intensity plots. The speed regulator is shown in grey. Right panel: Kymographs.

**Supplementary Figure 9. Analysis of hbA enhancer activity on a fixed embryo.** Images of a fixed hbA>MS2-*yellow*; aTub>MCP-mEmerlad *Tribolium* embryo in the germband stage (upper image: overview; lower image: close-up view of the indicated region in the overview image). The bright mEmerald puncta (MS2-MCP signal at active transcription sites) refine into a stripe resembling the *yellow* expression of the hbA enhancer reporter visualized using *in situ* HCR staining (compare to Figure 3E).

**Supplementary Figure 10. Visualizing runB>yellow expression waves in the germband using exonic vs intronic probes.** runB expression waves resemble more those of endogeneous *run* (shown in green) when visualized using intronic (shown in pruple) in situ probe than exonic (shown in red) probes.

**Supplementary Table 1:**
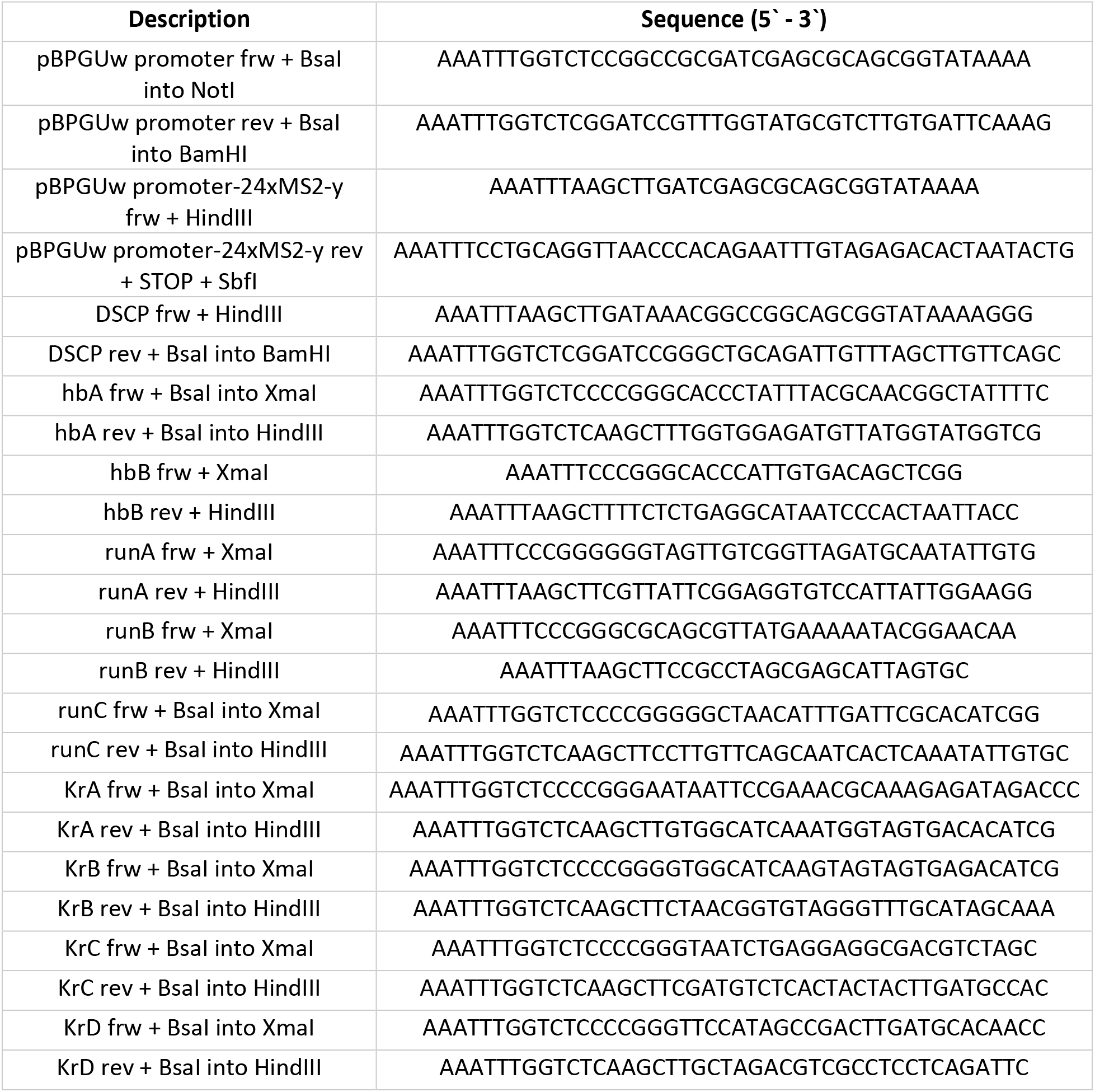
List of used primers. frw: forward, rev: reverse.

## Notes

### Competing Interest Statement

The authors have declared no competing interest.

### Summary of Updates

- Extended Discussion. - Adding simulations for enhancer deletion experiments (and associated Figure 7). - Updating some figures.

## References

1. Spitz F, Furlong EEM. Transcription factors: from enhancer binding to developmental control. Nat Rev Genet. 2012 Sep;13(9):613–626.

2. Shlyueva D, Stampfel G, Stark A. Transcriptional enhancers: from properties to genome-wide predictions. Nat Rev Genet. 2014 Apr;15(4):272–286.

3. Small S, Blair A, Levine M. Regulation of even-skipped stripe 2 in the Drosophila embryo. EMBO J. 1992 Nov;11(11):4047–4057.

4. Clyde DE, Corado MSG, Wu X, Paré A, Papatsenko D, Small S. A self-organizing system of repressor gradients establishes segmental complexity in Drosophila. Nature. 2003 Dec 18;426(6968):849–853.

5. Hoch M, Seifert E, Jäckle H. Gene expression mediated by cis-acting sequences of the Krüppel gene in response to the Drosophila morphogens bicoid and hunchback. EMBO J. 1991 Aug;10(8):2267–2278.

6. Clark E, Peel AD, Akam M. Arthropod segmentation. Development. 2019 Sep 25;146(18).

7. Lynch JA, El-Sherif E, Brown SJ. Comparisons of the embryonic development of Drosophila, Nasonia, and Tribolium. Wiley Interdiscip Rev Dev Biol. 2012 Feb;1(1):16–39.

8. Jaeger J. The gap gene network. Cell Mol Life Sci. 2011 Jan;68(2):243–274.

9. Diaz-Cuadros M, Pourquié O, El-Sherif E. Patterning with clocks and genetic cascades: Segmentation and regionalization of vertebrate versus insect body plans. PLoS Genet. 2021 Oct 14;17(10):e1009812.

10. Di Talia S, Vergassola M. Waves in embryonic development. Annu Rev Biophys. 2022 May 9;51:327–353.

11. Palmeirim I, Henrique D, Ish-Horowicz D, Pourquié O. Avian hairy gene expression identifies a molecular clock linked to vertebrate segmentation and somitogenesis. Cell. 1997 Nov 28;91(5):639–648.

12. Oates AC, Morelli LG, Ares S. Patterning embryos with oscillations: structure, function and dynamics of the vertebrate segmentation clock. Development. 2012 Feb;139(4):625–639.

13. Lauschke VM, Tsiairis CD, François P, Aulehla A. Scaling of embryonic patterning based on phase-gradient encoding. Nature. 2013 Jan 3;493(7430):101–105.

14. Aulehla A, Wiegraebe W, Baubet V, Wahl MB, Deng C, Taketo M, et al. A beta-catenin gradient links the clock and wavefront systems in mouse embryo segmentation. Nat Cell Biol. 2008 Feb 1;10(2):186–193.

15. Soroldoni D, Jörg DJ, Morelli LG, Richmond DL, Schindelin J, Jülicher F, et al. Genetic oscillations. A Doppler effect in embryonic pattern formation. Science. 2014 Jul 11;345(6193):222–225.

16. Sonnen KF, Janda CY. Signalling dynamics in embryonic development. Biochem J. 2021 Dec 10;478(23):4045–4070.

17. Forlani S, Lawson KA, Deschamps J. Acquisition of Hox codes during gastrulation and axial elongation in the mouse embryo. Development. 2003 Aug 1;130(16):3807–3819.

18. Gaunt SJ, Strachan L. Forward spreading in the establishment of a vertebrate Hox expression boundary: the expression domain separates into anterior and posterior zones, and the spread occurs across implanted glass barriers. Dev Dyn. 1994 Mar;199(3):229–240.

19. Deschamps J, Duboule D. Embryonic timing, axial stem cells, chromatin dynamics, and the Hox clock. Genes Dev. 2017 Jul 15;31(14):1406–1416.

20. Durston A, Wacker S, Bardine N, Jansen H. Time space translation: a hox mechanism for vertebrate a-p patterning. Curr Genomics. 2012 Jun;13(4):300–307.

21. Durston AJ, Jansen HJ, Wacker SA. Review: Time-space translation regulates trunk axial patterning in the early vertebrate embryo. Genomics. 2010 May;95(5):250–255.

22. Dessaud E, Yang LL, Hill K, Cox B, Ulloa F, Ribeiro A, et al. Interpretation of the sonic hedgehog morphogen gradient by a temporal adaptation mechanism. Nature. 2007 Nov 29;450(7170):717–720.

23. Balaskas N, Ribeiro A, Panovska J, Dessaud E, Sasai N, Page KM, et al. Gene regulatory logic for reading the Sonic Hedgehog signaling gradient in the vertebrate neural tube. Cell. 2012 Jan 20;148(1-2):273–284.

24. Harfe BD, Scherz PJ, Nissim S, Tian H, McMahon AP, Tabin CJ. Evidence for an expansion-based temporal Shh gradient in specifying vertebrate digit identities. Cell. 2004 Aug 20;118(4):517–528.

25. Saiz-Lopez P, Chinnaiya K, Campa VM, Delgado I, Ros MA, Towers M. An intrinsic timer specifies distal structures of the vertebrate limb. Nat Commun. 2015 Sep 18;6:8108.

26. Roensch K, Tazaki A, Chara O, Tanaka EM. Progressive specification rather than intercalation of segments during limb regeneration. Science. 2013 Dec 13;342(6164):1375–1379.

27. Towers M, Tickle C. Growing models of vertebrate limb development. Development. 2009 Jan;136(2):179–190.

28. El-Sherif E, Averof M, Brown SJ. A segmentation clock operating in blastoderm and germband stages of Tribolium development. Development. 2012 Dec 1;139(23):4341–4346.

29. Sarrazin AF, Peel AD, Averof M. A segmentation clock with two-segment periodicity in insects. Science. 2012 Apr 20;336(6079):338–341.

30. El-Sherif E, Zhu X, Fu J, Brown SJ. Caudal regulates the spatiotemporal dynamics of pair-rule waves in Tribolium. PLoS Genet. 2014 Oct 16;10(10):e1004677.

31. Zhu X, Rudolf H, Healey L, François P, Brown SJ, Klingler M, et al. Speed regulation of genetic cascades allows for evolvability in the body plan specification of insects. Proc Natl Acad Sci USA. 2017 Sep 25;114(41).

32. Boos A, Distler J, Rudolf H, Klingler M, El-Sherif E. A re-inducible gap gene cascade patterns the anterior-posterior axis of insects in a threshold-free fashion. Elife. 2018 Dec 20;7.

33. Brena C, Akam M. An analysis of segmentation dynamics throughout embryogenesis in the centipede Strigamia maritima. BMC Biol. 2013 Nov 29;11:112.

34. Pueyo JI, Lanfear R, Couso JP. Ancestral Notch-mediated segmentation revealed in the cockroach Periplaneta americana. Proc Natl Acad Sci USA. 2008 Oct 28;105(43):16614–16619.

35. Stollewerk A, Schoppmeier M, Damen WGM. Involvement of Notch and Delta genes in spider segmentation. Nature. 2003 Jun 19;423(6942):863–865.

36. Rosenberg MI, Brent AE, Payre F, Desplan C. Dual mode of embryonic development is highlighted by expression and function of Nasonia pair-rule genes. Elife. 2014 Mar 5;3(3):e01440.

37. Kanayama M, Akiyama-Oda Y, Nishimura O, Tarui H, Agata K, Oda H. Travelling and splitting of a wave of hedgehog expression involved in spider-head segmentation. Nat Commun. 2011 Oct 11;2:500.

38. Chipman AD, Akam M. The segmentation cascade in the centipede Strigamia maritima: involvement of the Notch pathway and pair-rule gene homologues. Dev Biol. 2008 Jul 1;319(1):160–169.

39. Pechmann M, McGregor AP, Schwager EE, Feitosa NM, Damen WGM. Dynamic gene expression is required for anterior regionalization in a spider. Proc Natl Acad Sci USA. 2009 Feb 3;106(5):1468–1472.

40. Jaeger J, Surkova S, Blagov M, Janssens H, Kosman D, Kozlov KN, et al. Dynamic control of positional information in the early Drosophila embryo. Nature. 2004 Jul 15;430(6997):368–371.

41. El-Sherif E, Levine M. Shadow enhancers mediate dynamic shifts of gap gene expression in the drosophila embryo. Curr Biol. 2016 May 9;26(9):1164–1169.

42. Lim B, Fukaya T, Heist T, Levine M. Temporal dynamics of pair-rule stripes in living Drosophila embryos. Proc Natl Acad Sci USA. 2018 Aug 14;115(33):8376–8381.

43. Berrocal A, Lammers NC, Garcia HG, Eisen MB. Kinetic sculpting of the seven stripes of the Drosophila even-skipped gene. Elife. 2020 Dec 10;9.

44. Keränen SVE, Fowlkes CC, Luengo Hendriks CL, Sudar D, Knowles DW, Malik J, et al. Three-dimensional morphology and gene expression in the Drosophila blastoderm at cellular resolution II: dynamics. Genome Biol. 2006;7(12):R124.

45. Rudolf H, Zellner C, El-Sherif E. Speeding up anterior-posterior patterning of insects by differential initialization of the gap gene cascade. Dev Biol. 2020 Apr 1;460(1):20–31.

46. Verd B, Clark E, Wotton KR, Janssens H, Jiménez-Guri E, Crombach A, et al. A damped oscillator imposes temporal order on posterior gap gene expression in Drosophila. PLoS Biol. 2018 Feb 16;16(2):e2003174.

47. Clark E, Peel AD. Evidence for the temporal regulation of insect segmentation by a conserved sequence of transcription factors. Development. 2018 May 3;

48. Clark E. Dynamic patterning by the Drosophila pair-rule network reconciles long-germ and short-germ segmentation. PLoS Biol. 2017 Sep 27;15(9):e2002439.

49. Kuhlmann L, El-Sherif E. Speed regulation and gradual enhancer switching models as flexible and evolvable patterning mechanisms. BioRxiv. 2018 Feb 7;

50. Falk HJ, Tomita T, Mönke G, McDole K, Aulehla A. Imaging the onset of oscillatory signaling dynamics during mouse embryo gastrulation. Development. 2022 Jul 1;149(13).

51. Diaz-Cuadros M, Wagner DE, Budjan C, Hubaud A, Tarazona OA, Donelly S, et al. In vitro characterization of the human segmentation clock. Nature. 2020 Apr;580(7801):113–118.

52. Shih NP, François P, Delaune EA, Amacher SL. Dynamics of the slowing segmentation clock reveal alternating two-segment periodicity. Development. 2015 May 15;142(10):1785–1793.

53. Hoermann A, Cicin-Sain D, Jaeger J. A quantitative validated model reveals two phases of transcriptional regulation for the gap gene giant in Drosophila. Dev Biol. 2016 Mar 15;411(2):325–338.

54. Oginuma M, Takahashi Y, Kitajima S, Kiso M, Kanno J, Kimura A, et al. The oscillation of Notch activation, but not its boundary, is required for somite border formation and rostral-caudal patterning within a somite. Development. 2010 May;137(9):1515–1522.

55. Shifley ET, Vanhorn KM, Perez-Balaguer A, Franklin JD, Weinstein M, Cole SE. Oscillatory lunatic fringe activity is crucial for segmentation of the anterior but not posterior skeleton. Development. 2008 Mar;135(5):899–908.

56. Stauber M, Sachidanandan C, Morgenstern C, Ish-Horowicz D. Differential axial requirements for lunatic fringe and Hes7 transcription during mouse somitogenesis. PLoS One. 2009 Nov 24;4(11):e7996.

57. Schroeder MD, Greer C, Gaul U. How to make stripes: deciphering the transition from non-periodic to periodic patterns in Drosophila segmentation. Development. 2011 Jul;138(14):3067–3078.

58. Schroeder MD, Pearce M, Fak J, Fan H, Unnerstall U, Emberly E, et al. Transcriptional control in the segmentation gene network of Drosophila. PLoS Biol. 2004 Sep;2(9):E271.

59. Bucher G, Scholten J, Klingler M. Parental rnai in tribolium (coleoptera). Curr Biol. 2002 Feb 5;12(3):R85–6.

60. Miller SC, Miyata K, Brown SJ, Tomoyasu Y. Dissecting systemic RNA interference in the red flour beetle Tribolium castaneum: parameters affecting the efficiency of RNAi. PLoS One. 2012 Oct 25;7(10):e47431.

61. Tomoyasu Y, Miller SC, Tomita S, Schoppmeier M, Grossmann D, Bucher G. Exploring systemic RNA interference in insects: a genome-wide survey for RNAi genes in Tribolium. Genome Biol. 2008 Jan 17;9(1):R10.

62. Dönitz J, Schmitt-Engel C, Grossmann D, Gerischer L, Tech M, Schoppmeier M, et al. iBeetle-Base: a database for RNAi phenotypes in the red flour beetle Tribolium castaneum. Nucleic Acids Res. 2015 Jan;43(Database issue):D720–5.

63. Choe CP, Brown SJ. Genetic regulation of engrailed and wingless in Tribolium segmentation and the evolution of pair-rule segmentation. Dev Biol. 2009 Jan 15;325(2):482–491.

64. Bolognesi R, Beermann A, Farzana L, Wittkopp N, Lutz R, Balavoine G, et al. Tribolium Wnts: evidence for a larger repertoire in insects with overlapping expression patterns that suggest multiple redundant functions in embryogenesis. Dev Genes Evol. 2008 Apr 8;218(3-4):193–202.

65. Choe CP, Miller SC, Brown SJ. A pair-rule gene circuit defines segments sequentially in the short-germ insect Tribolium castaneum. Proc Natl Acad Sci USA. 2006 Apr 25;103(17):6560–6564.

66. Kotkamp K, Klingler M, Schoppmeier M. Apparent role of Tribolium orthodenticle in anteroposterior blastoderm patterning largely reflects novel functions in dorsoventral axis formation and cell survival. Development. 2010 Jun;137(11):1853–1862.

67. Schoppmeier M, Fischer S, Schmitt-Engel C, Löhr U, Klingler M. An ancient anterior patterning system promotes caudal repression and head formation in ecdysozoa. Curr Biol. 2009 Nov 17;19(21):1811–1815.

68. Cerny AC, Bucher G, Schröder R, Klingler M. Breakdown of abdominal patterning in the Tribolium Kruppel mutant jaws. Development. 2005 Dec;132(24):5353–5363.

69. Bucher G, Klingler M. Divergent segmentation mechanism in the short germ insect Tribolium revealed by giant expression and function. Development. 2004 Apr;131(8):1729–1740.

70. Schmitt-Engel C, Cerny AC, Schoppmeier M. A dual role for nanos and pumilio in anterior and posterior blastodermal patterning of the short-germ beetle Tribolium castaneum. Dev Biol. 2012 Apr 15;364(2):224–235.

71. Marques-Souza H, Aranda M, Tautz D. Delimiting the conserved features of hunchback function for the trunk organization of insects. Development. 2008 Mar;135(5):881–888.

72. Savard J, Marques-Souza H, Aranda M, Tautz D. A segmentation gene in tribolium produces a polycistronic mRNA that codes for multiple conserved peptides. Cell. 2006 Aug 11;126(3):559–569.

73. Marques-Souza H. Evolution of the gene regulatory network controlling trunk segmentation in insects [Doctoral dissertation]. University of Cologne; 2007.

74. Jeon H, Gim S, Na H, Choe CP. A pair-rule function of odd-skipped in germband stages of Tribolium development. Dev Biol. 2020 Jul 17;

75. Klingler M, Bucher G. The red flour beetle T. castaneum: elaborate genetic toolkit and unbiased large scale RNAi screening to study insect biology and evolution. Evodevo. 2022 Dec;13(1):14.

76. Pavlopoulos A, Berghammer AJ, Averof M, Klingler M. Efficient transformation of the beetle Tribolium castaneum using the Minos transposable element: quantitative and qualitative analysis of genomic integration events. Genetics. 2004 Jun;167(2):737–746.

77. Strobl F, Anderl A, Stelzer EH. A universal vector concept for a direct genotyping of transgenic organisms and a systematic creation of homozygous lines. Elife. 2018 Mar 15;7.

78. Trauner J, Schinko J, Lorenzen MD, Shippy TD, Wimmer EA, Beeman RW, et al. Large-scale insertional mutagenesis of a coleopteran stored grain pest, the red flour beetle Tribolium castaneum, identifies embryonic lethal mutations and enhancer traps. BMC Biol. 2009 Nov 5;7:73.

79. Gilles AF, Schinko JB, Averof M. Efficient CRISPR-mediated gene targeting and transgene replacement in the beetle Tribolium castaneum. Development. 2015 Aug 15;142(16):2832–2839.

80. Gilles AF, Schinko JB, Schacht MI, Enjolras C, Averof M. Clonal analysis by tunable CRISPR-mediated excision. Development. 2019 Jan 4;146(1).

81. Strobl F, Stelzer EH. Long-term fluorescence live imaging of Tribolium castaneum embryos: principles, resources, scientific challenges and the comparative approach. Curr Opin Insect Sci. 2016 Aug 28;18:17–26.

82. Strobl F, Stelzer EHK. Non-invasive long-term fluorescence live imaging of Tribolium castaneum embryos. Development. 2014 Jun;141(11):2331–2338.

83. Strobl F, Schmitz A, Stelzer EHK. Live imaging of Tribolium castaneum embryonic development using light-sheet-based fluorescence microscopy. Nat Protoc. 2015 Oct;10(10):1486–1507.

84. Macaya CC, Saavedra PE, Cepeda RE, Nuñez VA, Sarrazin AF. A Tribolium castaneum whole embryo culture protocol for studying the molecular mechanisms and morphogenetic movements involved in insect development. Dev Genes Evol. 2016 Jan 6;226(1):53–61.

85. Buenrostro JD, Giresi PG, Zaba LC, Chang HY, Greenleaf WJ. Transposition of native chromatin for fast and sensitive epigenomic profiling of open chromatin, DNA-binding proteins and nucleosome position. Nat Methods. 2013 Dec;10(12):1213–1218.

86. Thurman RE, Rynes E, Humbert R, Vierstra J, Maurano MT, Haugen E, et al. The accessible chromatin landscape of the human genome. Nature. 2012 Sep 6;489(7414):75–82.

87. Kok K, Arnosti DN. Dynamic reprogramming of chromatin: paradigmatic palimpsests and HES factors. Front Genet. 2015 Feb 10;6:29.

88. Xi H, Shulha HP, Lin JM, Vales TR, Fu Y, Bodine DM, et al. Identification and characterization of cell type-specific and ubiquitous chromatin regulatory structures in the human genome. PLoS Genet. 2007 Aug;3(8):e136.

89. Li LM, Arnosti DN. Long-and short-range transcriptional repressors induce distinct chromatin states on repressed genes. Curr Biol. 2011 Mar 8;21(5):406–412.

90. Reddington JP, Garfield DA, Sigalova OM, Karabacak Calviello A, Marco-Ferreres R, Girardot C, et al. Lineage-Resolved Enhancer and Promoter Usage during a Time Course of Embryogenesis. Dev Cell. 2020 Dec 7;55(5):648–664.e9.

91. Bozek M, Cortini R, Storti AE, Unnerstall U, Gaul U, Gompel N. ATAC-seq reveals regional differences in enhancer accessibility during the establishment of spatial coordinates in the Drosophila blastoderm. Genome Res. 2019 May;29(5):771–783.

92. McKay DJ, Lieb JD. A common set of DNA regulatory elements shapes Drosophila appendages. Dev Cell. 2013 Nov 11;27(3):306–318.

93. Li X, Zhao X, Fang Y, Jiang X, Duong T, Fan C, et al. Generation of destabilized green fluorescent protein as a transcription reporter. J Biol Chem. 1998 Dec 25;273(52):34970–34975.

94. He L, Binari R, Huang J, Falo-Sanjuan J, Perrimon N. In vivo study of gene expression with an enhanced dual-color fluorescent transcriptional timer. Elife. 2019 May 29;8.

95. Pichon X, Lagha M, Mueller F, Bertrand E. A growing toolbox to image gene expression in single cells: sensitive approaches for demanding challenges. Mol Cell. 2018 Aug 2;71(3):468–480.

96. Johansson HE, Liljas L, Uhlenbeck OC. RNA recognition by the MS2 phage coat protein. Seminars in virology. 1997;8(3):176–185.

97. Garcia HG, Tikhonov M, Lin A, Gregor T. Quantitative imaging of transcription in living Drosophila embryos links polymerase activity to patterning. Curr Biol. 2013 Nov 4;23(21):2140–2145.

98. Lucas T, Ferraro T, Roelens B, De Las Heras Chanes J, Walczak AM, Coppey M, et al. Live imaging of bicoid-dependent transcription in Drosophila embryos. Curr Biol. 2013 Nov 4;23(21):2135–2139.

99. Bothma JP, Garcia HG, Esposito E, Schlissel G, Gregor T, Levine M. Dynamic regulation of eve stripe 2 expression reveals transcriptional bursts in living Drosophila embryos. Proc Natl Acad Sci USA. 2014 Jul 22;111(29):10598–10603.

100. Simon JM, Giresi PG, Davis IJ, Lieb JD. Using formaldehyde-assisted isolation of regulatory elements (FAIRE) to isolate active regulatory DNA. Nat Protoc. 2012 Jan 19;7(2):256–267.

101. Lai Y-T, Deem KD, Borràs-Castells F, Sambrani N, Rudolf H, Suryamohan K, et al. Enhancer identification and activity evaluation in the red flour beetle, Tribolium castaneum. Development. 2018 Apr 5;145(7).

102. Perry MW, Boettiger AN, Levine M. Multiple enhancers ensure precision of gap gene-expression patterns in the Drosophila embryo. Proc Natl Acad Sci USA. 2011 Aug 16;108(33):13570–13575.

103. Choi HMT, Schwarzkopf M, Fornace ME, Acharya A, Artavanis G, Stegmaier J, et al. Third-generation in situ hybridization chain reaction: multiplexed, quantitative, sensitive, versatile, robust. Development. 2018 Jun 26;145(12).

104. Cande J, Goltsev Y, Levine MS. Conservation of enhancer location in divergent insects. Proc Natl Acad Sci USA. 2009 Aug 25;106(34):14414–14419.

105. Schönauer A, Paese CLB, Hilbrant M, Leite DJ, Schwager EE, Feitosa NM, et al. The Wnt and Delta-Notch signalling pathways interact to direct pair-rule gene expression via caudal during segment addition in the spider Parasteatoda tepidariorum. Development. 2016 Jul 1;143(13):2455–2463.

106. Bolognesi R, Farzana L, Fischer TD, Brown SJ. Multiple Wnt genes are required for segmentation in the short-germ embryo of Tribolium castaneum. Curr Biol. 2008 Oct 28;18(20):1624–1629.

107. Copf T, Schröder R, Averof M. Ancestral role of caudal genes in axis elongation and segmentation. Proc Natl Acad Sci USA. 2004 Dec 21;101(51):17711–17715.

108. McGregor AP, Pechmann M, Schwager EE, Feitosa NM, Kruck S, Aranda M, et al. Wnt8 is required for growth-zone establishment and development of opisthosomal segments in a spider. Curr Biol. 2008 Oct 28;18(20):1619–1623.

109. Bailles A, Gehrels EW, Lecuit T. Mechanochemical principles of spatial and temporal patterns in cells and tissues. Annu Rev Cell Dev Biol. 2022 May 13;

110. Jouve C, Iimura T, Pourquie O. Onset of the segmentation clock in the chick embryo: evidence for oscillations in the somite precursors in the primitive streak. Development. 2002 Mar;129(5):1107–1117.

111. Jutras-Dubé L, El-Sherif E, François P. Geometric models for robust encoding of dynamical information into embryonic patterns. Elife. 2020 Aug 10;9.

112. Vroomans RMA, Hogeweg P, Ten Tusscher KHWJ. Around the clock: gradient shape and noise impact the evolution of oscillatory segmentation dynamics. Evodevo. 2018 Dec 10;9:24.

113. Winfree AT. The Geometry of Biological Time (Interdisciplinary Applied Mathematics). 2nd ed. 2001. Softcover reprint of the original 2nd ed. 2001. Springer; 2010.

114. Rayon T, Stamataki D, Perez-Carrasco R, Garcia-Perez L, Barrington C, Melchionda M, et al. Species-specific pace of development is associated with differences in protein stability. Science. 2020 Sep 18;369(6510).

115. Uyehara CM, Nystrom SL, Niederhuber MJ, Leatham-Jensen M, Ma Y, Buttitta LA, et al. Hormone-dependent control of developmental timing through regulation of chromatin accessibility. Genes Dev. 2017 May 1;31(9):862–875.

116. Pease NA, Nguyen PHB, Woodworth MA, Ng KKH, Irwin B, Vaughan JC, et al. Tunable, division-independent control of gene activation timing by a polycomb switch. Cell Rep. 2021 Mar 23;34(12):108888.

117. Nguyen P, Pease NA, Kueh HY. Scalable control of developmental timetables by epigenetic switching networks. J R Soc Interface. 2021 Jul 21;18(180):20210109.

118. Benito-Kwiecinski S, Giandomenico SL, Sutcliffe M, Riis ES, Freire-Pritchett P, Kelava I, et al. An early cell shape transition drives evolutionary expansion of the human forebrain. Cell. 2021 Apr 15;184(8):2084–2102.e19.

119. Bielen H, Pal S, Tole S, Houart C. Temporal variations in early developmental decisions: an engine of forebrain evolution. Curr Opin Neurobiol. 2017 Feb;42:152–159.

120. Preger-Ben Noon E, Sabarís G, Ortiz DM, Sager J, Liebowitz A, Stern DL, et al. Comprehensive Analysis of a cis-Regulatory Region Reveals Pleiotropy in Enhancer Function. Cell Rep. 2018 Mar 13;22(11):3021–3031.

121. Sabarís G, Laiker I, Preger-Ben Noon E, Frankel N. Actors with Multiple Roles: Pleiotropic Enhancers and the Paradigm of Enhancer Modularity. Trends Genet. 2019 Jun;35(6):423–433.

122. Lewis JJ, Geltman RC, Pollak PC, Rondem KE, Van Belleghem SM, Hubisz MJ, et al. Parallel evolution of ancient, pleiotropic enhancers underlies butterfly wing pattern mimicry. Proc Natl Acad Sci USA. 2019 Nov 26;116(48):24174–24183.

123. Murugesan SN, Connahs H, Matsuoka Y, Das Gupta M, Tiong GJL, Huq M, et al. Butterfly eyespots evolved via cooption of an ancestral gene-regulatory network that also patterns antennae, legs, and wings. Proc Natl Acad Sci USA. 2022 Feb 22;119(8).

124. Erceg J, Pakozdi T, Marco-Ferreres R, Ghavi-Helm Y, Girardot C, Bracken AP, et al. Dual functionality of cis-regulatory elements as developmental enhancers and Polycomb response elements. Genes Dev. 2017 Mar 15;31(6):590–602.

125. Arnosti DN, Barolo S, Levine M, Small S. The eve stripe 2 enhancer employs multiple modes of transcriptional synergy. Development. 1996 Jan;122(1):205–214.

126. Howard K, Ingham P, Rushlow C. Region-specific alleles of the Drosophila segmentation gene hairy. Genes Dev. 1988 Aug;2(8):1037–1046.

127. Estella C, Voutev R, Mann RS. A dynamic network of morphogens and transcription factors patterns the fly leg. Curr Top Dev Biol. 2012;98:173–198.

128. Clark E, Akam M. Odd-paired controls frequency doubling in Drosophila segmentation by altering the pair-rule gene regulatory network. Elife. 2016 Aug 15;5.

129. Soluri IV, Zumerling LM, Payan Parra OA, Clark EG, Blythe SA. Zygotic pioneer factor activity of Odd-paired/Zic is necessary for late function of the Drosophila segmentation network. Elife. 2020 Apr 29;9.

130. Koromila T, Gao F, Iwasaki Y, He P, Pachter L, Gergen JP, et al. Odd-paired is a pioneer-like factor that coordinates with Zelda to control gene expression in embryos. Elife. 2020 Jul 23;9.

131. Taylor SE, Dearden PK. The Nasonia pair-rule gene regulatory network retains its function over 300 million years of evolution. Development. 2022 Mar 1;149(5).

132. Ribeiro L, Tobias-Santos V, Santos D, Antunes F, Feltran G, de Souza Menezes J, et al. Evolution and multiple roles of the Pancrustacea specific transcription factor zelda in insects. PLoS Genet. 2017 Jul 3;13(7):e1006868.

133. Ng KK, Yui MA, Mehta A, Siu S, Irwin B, Pease S, et al. A stochastic epigenetic switch controls the dynamics of T-cell lineage commitment. Elife. 2018 Nov 20;7.

134. Ay A, Holland J, Sperlea A, Devakanmalai GS, Knierer S, Sangervasi S, et al. Spatial gradients of protein-level time delays set the pace of the traveling segmentation clock waves. Development. 2014 Nov;141(21):4158–4167.

135. Okubo Y, Sugawara T, Abe-Koduka N, Kanno J, Kimura A, Saga Y. Lfng regulates the synchronized oscillation of the mouse segmentation clock via trans-repression of Notch signalling. Nat Commun. 2012;3:1141.

136. Tsiairis CD, Aulehla A. Self-Organization of Embryonic Genetic Oscillators into Spatiotemporal Wave Patterns. Cell. 2016 Feb 11;164(4):656–667.

137. Herrgen L, Ares S, Morelli LG, Schröter C, Jülicher F, Oates AC. Intercellular coupling regulates the period of the segmentation clock. Curr Biol. 2010 Jul 27;20(14):1244–1253.

138. Yoshioka-Kobayashi K, Matsumiya M, Niino Y, Isomura A, Kori H, Miyawaki A, et al. Coupling delay controls synchronized oscillation in the segmentation clock. Nature. 2020 Apr;580(7801):119–123.

139. Morelli LG, Ares S, Herrgen L, Schröter C, Jülicher F, Oates AC. Delayed coupling theory of vertebrate segmentation. HFSP J. 2009;3(1):55–66.

140. Liao B-K, Jörg DJ, Oates AC. Faster embryonic segmentation through elevated Delta-Notch signalling. Nat Commun. 2016 Jun 15;7:11861.

141. Dönitz J, Gerischer L, Hahnke S, Pfeiffer S, Bucher G. Expanded and updated data and a query pipeline for iBeetle-Base. Nucleic Acids Res. 2018 Jan 4;46(D1):D831–D835.

142. Pfeiffer BD, Jenett A, Hammonds AS, Ngo T-TB, Misra S, Murphy C, et al. Tools for neuroanatomy and neurogenetics in Drosophila. Proc Natl Acad Sci USA. 2008 Jul 15;105(28):9715–9720.

143. Tribolium Genome Sequencing Consortium, Richards S, Gibbs RA, Weinstock GM, Brown SJ, Denell R, et al. The genome of the model beetle and pest Tribolium castaneum. Nature. 2008 Apr 24;452(7190):949–955.

144. Siebert KS, Lorenzen MD, Brown SJ, Park Y, Beeman RW. Tubulin superfamily genes in Tribolium castaneum and the use of a Tubulin promoter to drive transgene expression. Insect Biochem Mol Biol. 2008 Aug;38(8):749–755.

145. Chook YM, Blobel G. Karyopherins and nuclear import. Curr Opin Struct Biol. 2001 Dec;11(6):703–715.

146. Field J, Nikawa J, Broek D, MacDonald B, Rodgers L, Wilson IA, et al. Purification of a RAS-responsive adenylyl cyclase complex from Saccharomyces cerevisiae by use of an epitope addition method. Mol Cell Biol. 1988 May;8(5):2159–2165.

147. Bertrand E, Chartrand P, Schaefer M, Shenoy SM, Singer RH, Long RM. Localization of ASH1 mRNA particles in living yeast. Mol Cell. 1998 Oct;2(4):437–445.

148. Shaner NC, Steinbach PA, Tsien RY. A guide to choosing fluorescent proteins. Nat Methods. 2005 Dec;2(12):905–909.

149. Li X, Harrell RA, Handler AM, Beam T, Hennessy K, Fraser MJ. piggyBac internal sequences are necessary for efficient transformation of target genomes. Insect Mol Biol. 2005 Jan;14(1):17–30.

150. Shaner NC, Lin MZ, McKeown MR, Steinbach PA, Hazelwood KL, Davidson MW, et al. Improving the photostability of bright monomeric orange and red fluorescent proteins. Nat Methods. 2008 Jun;5(6):545–551.

151. Shaner NC, Campbell RE, Steinbach PA, Giepmans BNG, Palmer AE, Tsien RY. Improved monomeric red, orange and yellow fluorescent proteins derived from Discosoma sp. red fluorescent protein. Nat Biotechnol. 2004 Dec;22(12):1567–1572.

152. Berghammer AJ, Klingler M, Wimmer EA. A universal marker for transgenic insects. Nature. 1999 Nov 25;402(6760):370–371.

153. Handler AM, Harrell RA. Germline transformation of Drosophila melanogaster with the piggyBac transposon vector. Insect Mol Biol. 1999 Nov;8(4):449–457.

154. Lorenzen MD, Berghammer AJ, Brown SJ, Denell RE, Klingler M, Beeman RW. piggyBac-mediated germline transformation in the beetle Tribolium castaneum. Insect Mol Biol. 2003 Oct;12(5):433–440.

155. Berghammer AJ, Weber M, Trauner J, Klingler M. Red flour beetle (Tribolium) germline transformation and insertional mutagenesis. Cold Spring Harb Protoc. 2009 Aug;2009(8):pdb.prot5259.

156. Lorenzen MD, Brown SJ, Denell RE, Beeman RW. Cloning and characterization of the Tribolium castaneum eye-color genes encoding tryptophan oxygenase and kynurenine 3-monooxygenase. Genetics. 2002 Jan;160(1):225–234.

157. Handler AM. Use of the piggyBac transposon for germ-line transformation of insects. Insect Biochem Mol Biol. 2002 Oct;32(10):1211–1220.

158. Schindelin J, Arganda-Carreras I, Frise E, Kaynig V, Longair M, Pietzsch T, et al. Fiji: an open-source platform for biological-image analysis. Nat Methods. 2012 Jun 28;9(7):676–682.

159. Buenrostro JD, Wu B, Chang HY, Greenleaf WJ. ATAC-seq: a method for assaying chromatin accessibility genome-wide. Curr Protoc Mol Biol. 2015 Jan 5;109:21.29.1-21.29.9.

160. Blythe SA, Wieschaus EF. Establishment and maintenance of heritable chromatin structure during early Drosophila embryogenesis. Elife. 2016 Nov 23;5.

161. Martin M. Cutadapt removes adapter sequences from high-throughput sequencing reads. EMBnet j. 2011 May 2;17(1):10.

162. Aligning sequence reads, clone sequences and assembly contigs with BWA-MEM – ScienceOpen [Internet]. [cited 2022 Oct 24]. Available from: https://www.scienceopen.com/document?vid=e623e045-f570-42c5-80c8-ef0aea06629c

163. Danecek P, Bonfield JK, Liddle J, Marshall J, Ohan V, Pollard MO, et al. Twelve years of SAMtools and BCFtools. Gigascience. 2021 Feb 16;10(2).

164. Amemiya HM, Kundaje A, Boyle AP. The ENCODE blacklist: identification of problematic regions of the genome. Sci Rep. 2019 Jun 27;9(1):9354.

165. Pongor LS, Gross JM, Vera Alvarez R, Murai J, Jang S-M, Zhang H, et al. BAMscale: quantification of next-generation sequencing peaks and generation of scaled coverage tracks. Epigenetics Chromatin. 2020 Apr 22;13(1):21.

166. Zhang Y, Liu T, Meyer CA, Eeckhoute J, Johnson DS, Bernstein BE, et al. Model-based analysis of ChIP-Seq (MACS). Genome Biol. 2008 Sep 17;9(9):R137.

167. Heinz S, Benner C, Spann N, Bertolino E, Lin YC, Laslo P, et al. Simple combinations of lineage determining transcription factors prime cis-regulatory elements required for macrophage and B cell identities. Mol Cell. 2010 May 28;38(4):576–589.

168. Lopez-Delisle L, Rabbani L, Wolff J, Bhardwaj V, Backofen R, Grüning B, et al. pyGenomeTracks: reproducible plots for multivariate genomic datasets. Bioinformatics. 2021 Apr 20;37(3):422–423.

169. Ramírez F, Bhardwaj V, Arrigoni L, Lam KC, Grüning BA, Villaveces J, et al. High-resolution TADs reveal DNA sequences underlying genome organization in flies. Nat Commun. 2018 Jan 15;9(1):189.

170. Robinson MD, McCarthy DJ, Smyth GK. edgeR: a Bioconductor package for differential expression analysis of digital gene expression data. Bioinformatics. 2010 Jan 1;26(1):139–140.

171. Love MI, Huber W, Anders S. Moderated estimation of fold change and’ ‘ dispersion for RNA-seq data with DESeq2. Genome Biol. 2014;15(12):550.

172. Ross-Innes CS, Stark R, Teschendorff AE, Holmes KA, Ali HR, Dunning MJ, et al. Differential oestrogen receptor binding is associated with clinical outcome in breast cancer. Nature. 2012 Jan 4;481(7381):389–393.

173. Robinson MD, Oshlack A. A scaling normalization method for differential expression analysis of RNA-seq data. Genome Biol. 2010 Mar 2;11(3):R25.

174. Grant CE, Bailey TL. XSTREME: Comprehensive motif analysis of biological sequence datasets. BioRxiv. 2021 Sep 3;

175. Castro-Mondragon JA, Riudavets-Puig R, Rauluseviciute I, Lemma RB, Turchi L, Blanc-Mathieu R, et al. JASPAR 2022: the 9th release of the open-access database of transcription factor binding profiles. Nucleic Acids Res. 2022 Jan 7;50(D1):D165–D173.

