## Supplementary Figures for "How enhancers regulate wavelike gene expression patterns: Novel enhancer prediction and live reporter systems identify an enhancer associated with the arrest of pair-rule waves in the short-germ beetle *Tribolium*"

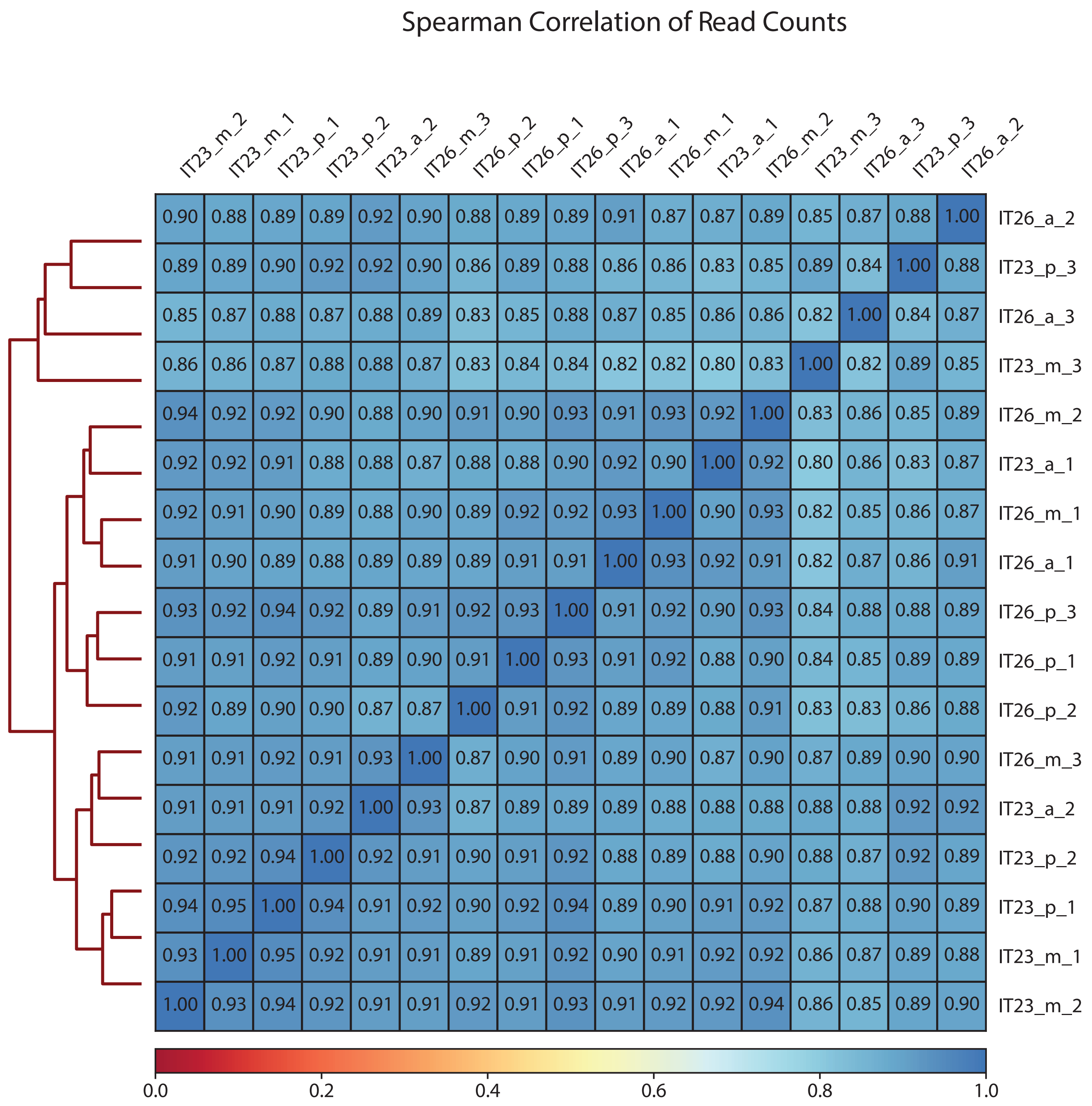


**Supplementary Figure 1: Correlation between ATAC-seq sequencing libraries.**

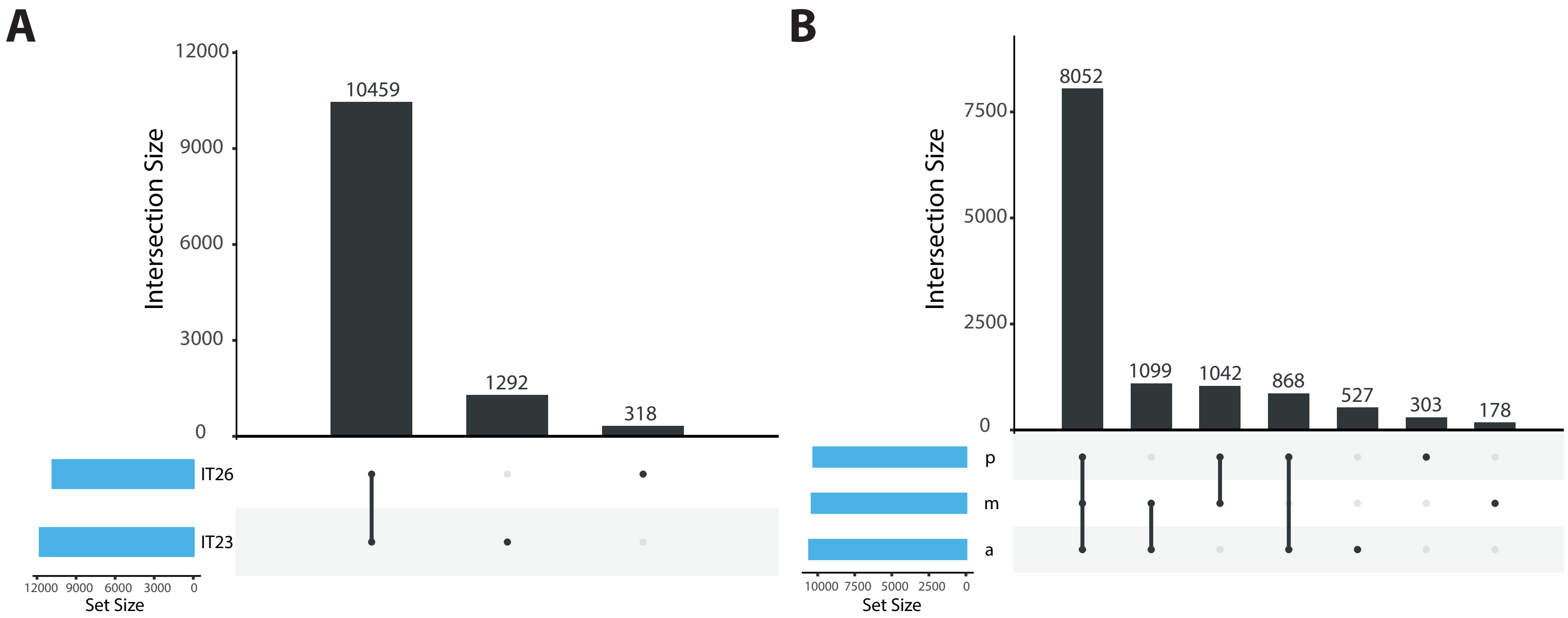


**Supplementary Figure 2: Upset plot comparing the number of sites identified in samples corresponding to IT23 and IT26 as well as along the AP axis.**

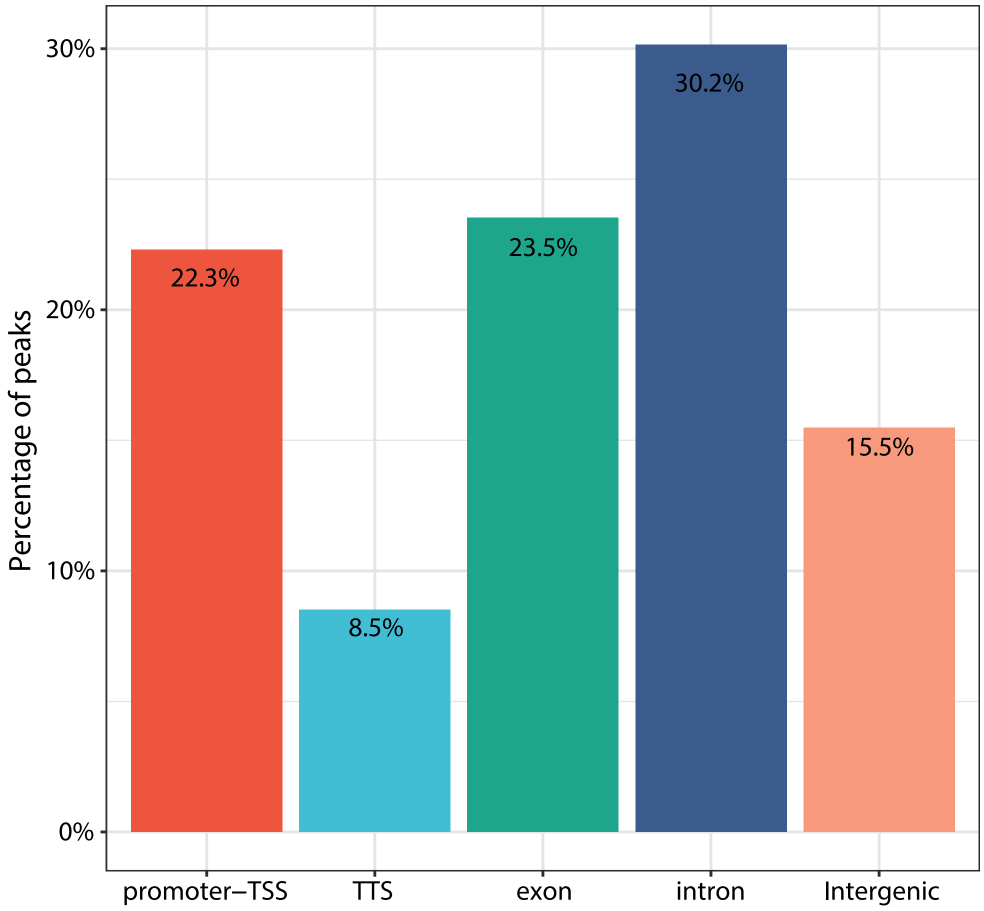


**Supplementary Figure 3: Annotation of consensus sites.**

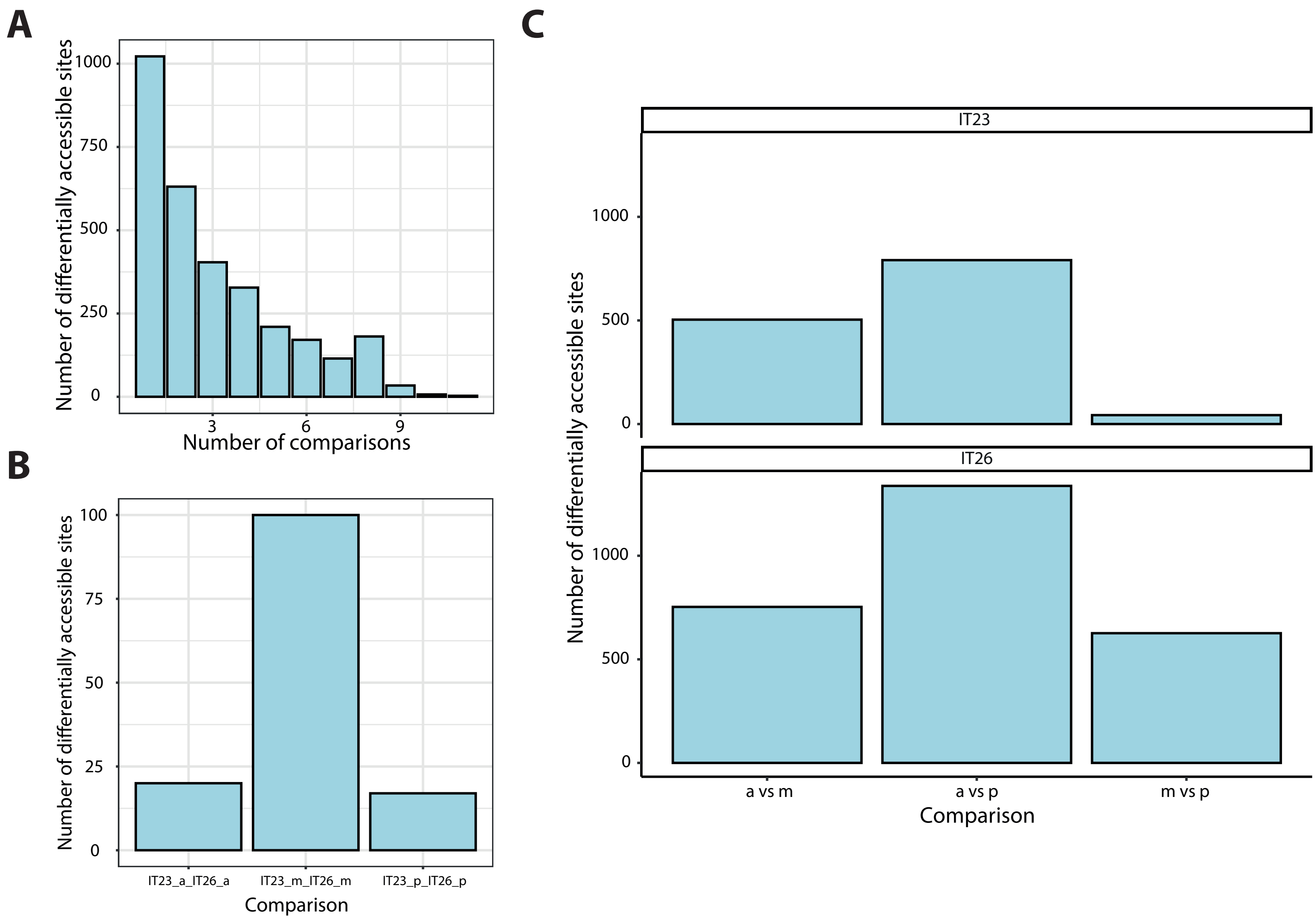


**Supplementary Figure 4: Analysis of differentially accessible sites.**

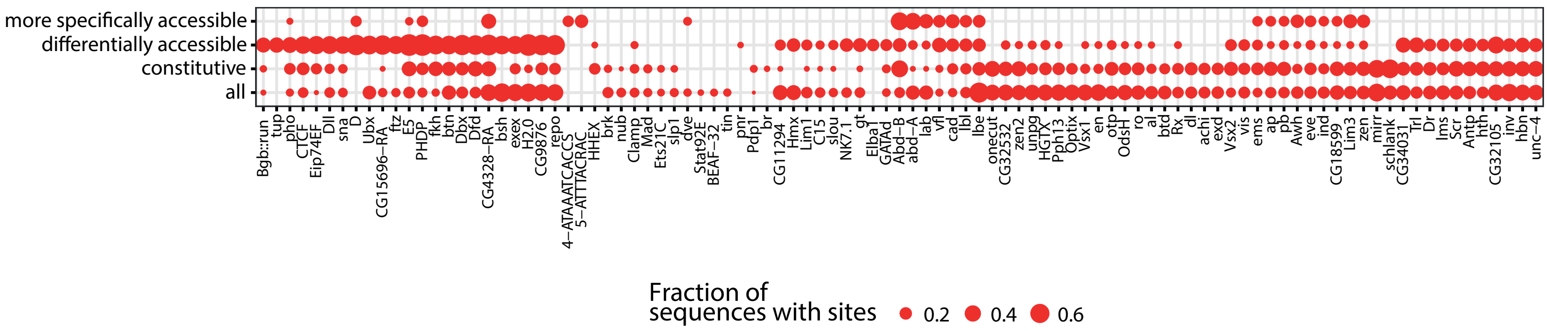


**Supplementary Figure 5: Motif Analysis.**

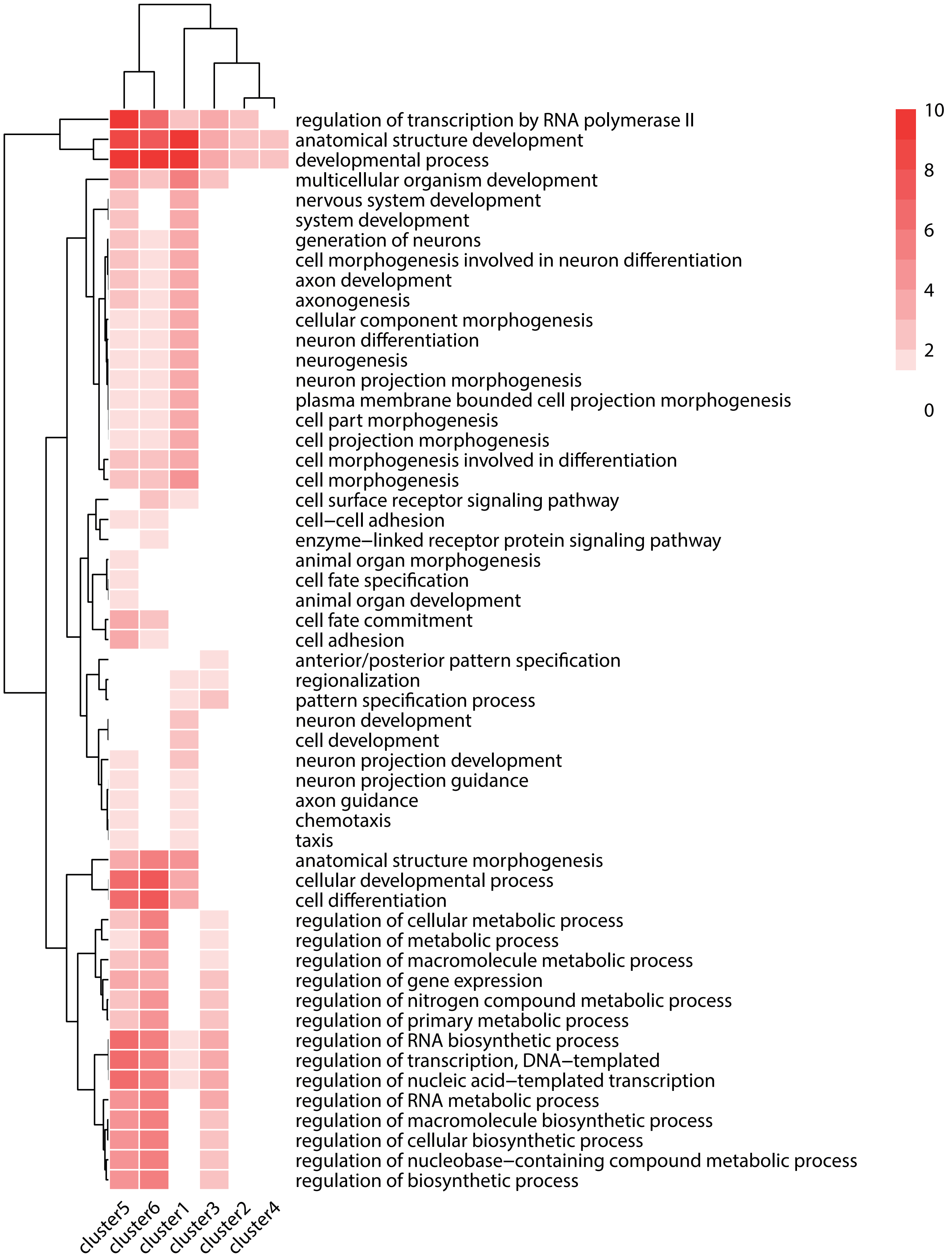


**Supplementary Figure 6: Functional enrichment analysis of the clusters of differentially accessible sites.**

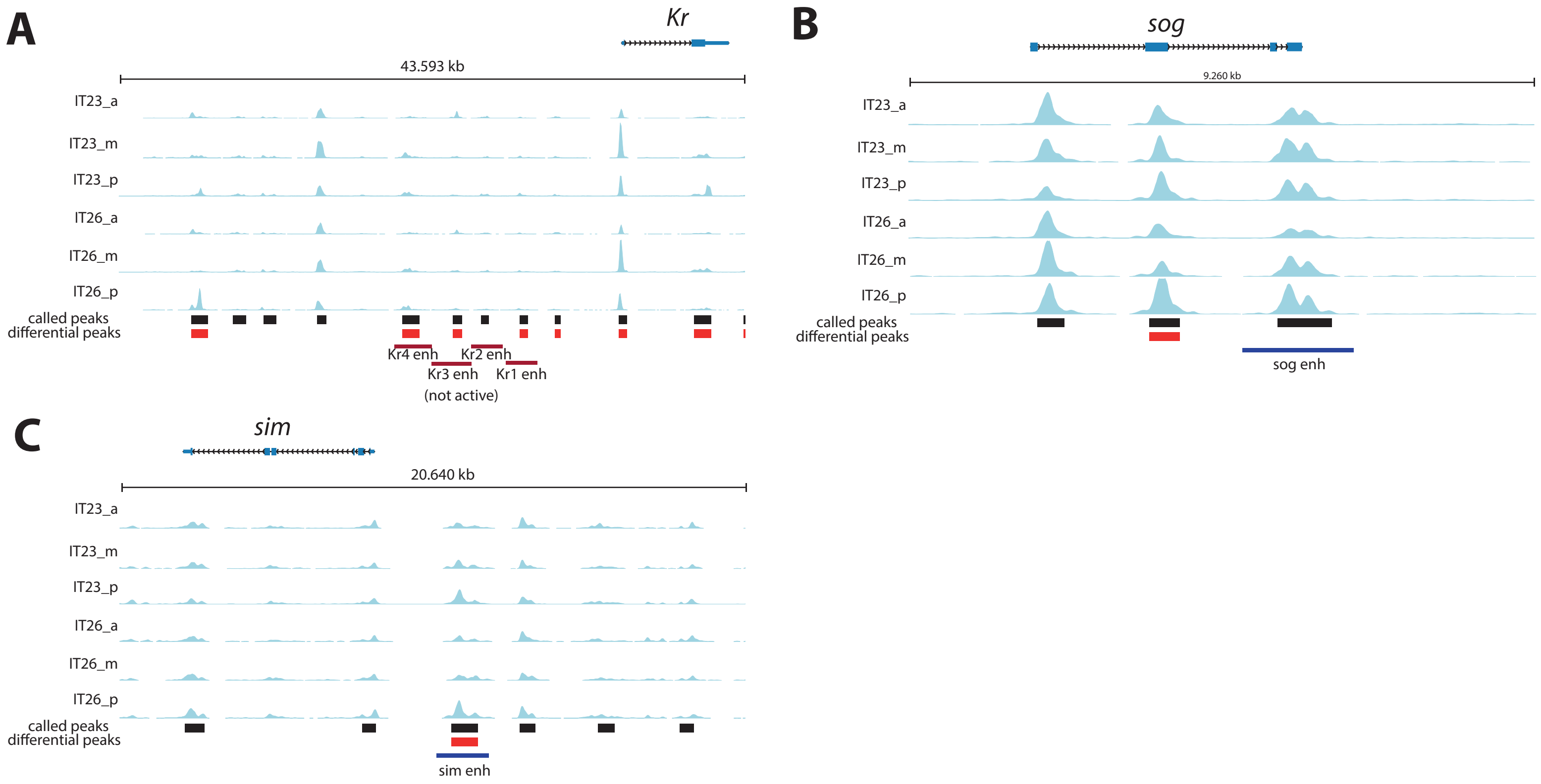


**Supplementary Figure 7: Genomic tracks of analyzed enhancer reporter constructs.**

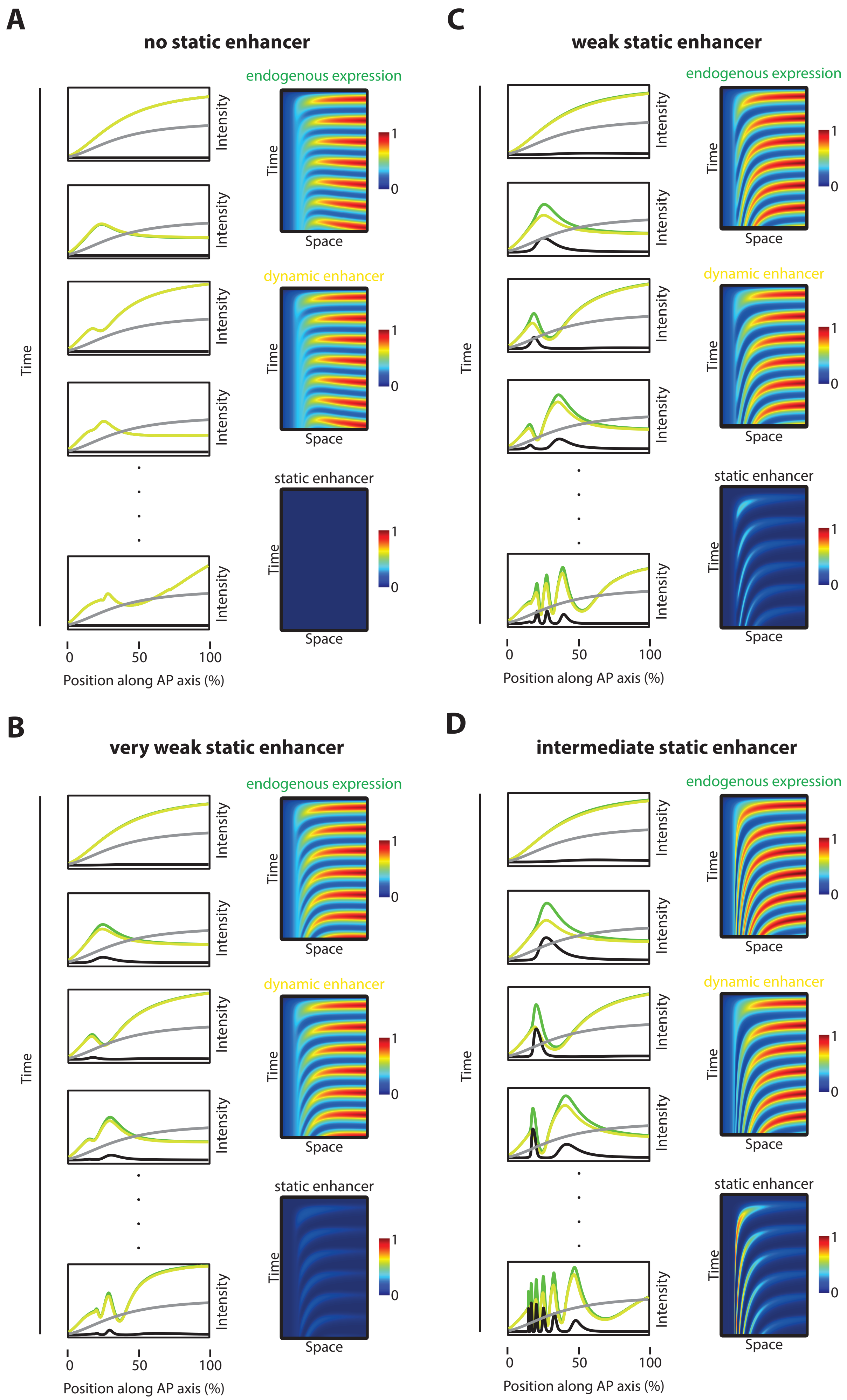


**Supplementary Figure 8: Simulations with different static enhancer strength.**

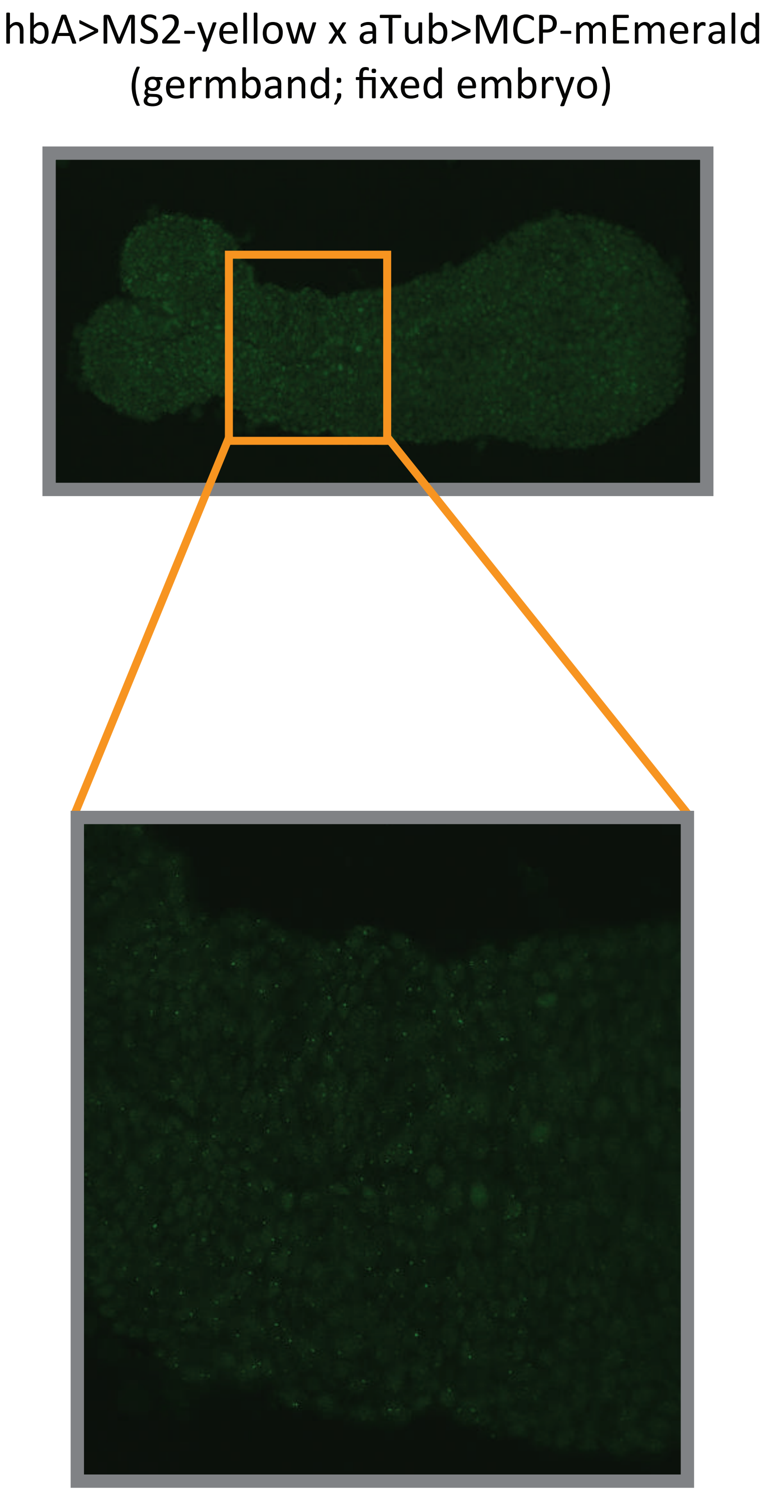


**Supplementary Figure 9: Analysis of hbA enhancer activity on a fixed embryo.**

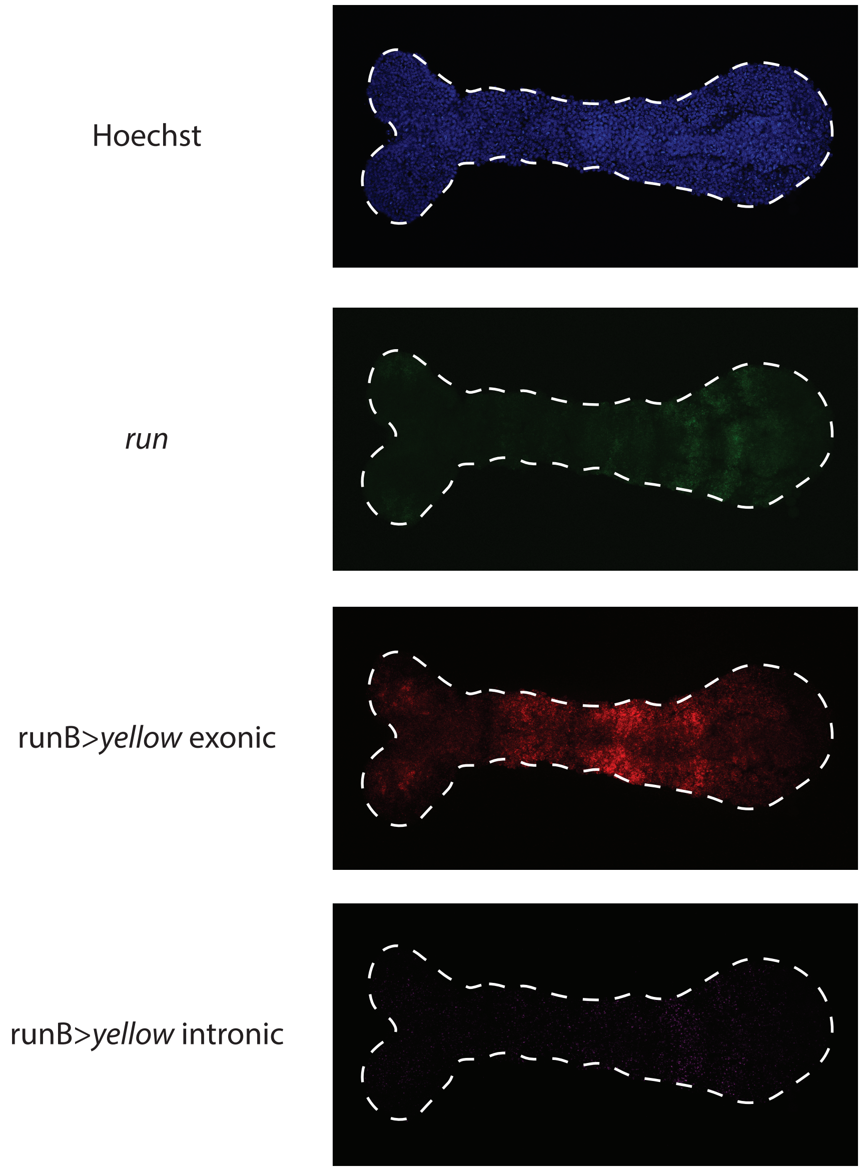


**Supplementary Figure 10: Visualizing runB>yellow expression waves in the germband using exonic vs intronic probes.**
